# Mechanism-based prediction of insertion-driven high pathogenicity avian influenza virus emergence

**DOI:** 10.64898/2026.08.27.747464

**Authors:** Gabriel Dupré, Bertille Pouget, Aldair Martinez-Pineda, Charlotte Foret-Lucas, Pierre Bessière, Delphine Chrétien, Mariette Ducatez, Nathalie Vialaneix, Claire Hoede, Roland Marquet, Christine Gaspin, Romain Volmer

## Abstract

High pathogenicity avian influenza viruses (HPAIVs) emerge from H5 and H7 low-pathogenicity avian influenza virus progenitors through mutations that introduce a multibasic cleavage site in haemagglutinin. Although nucleotide insertions recurrently generate this motif, the molecular determinants of insertion and whether particular HA sequences are genetically predisposed to evolve toward HPAIV remain unknown. Combining experimental virology and thermodynamic modelling, we show that insertions arise through polymerase slippage controlled by local product–template duplex thermodynamics within the viral polymerase catalytic site. Predicted RNA secondary structures outside the polymerase are not required for high-frequency insertions and only modestly modulate insertion rates. We formalize this mechanism in HPAIVpredict, which predicts insertion profiles, recapitulates intermediates associated with documented HPAIV emergence events and identifies H5 and H7 sequence backgrounds predisposed to acquire functional multibasic cleavage sites.

---

Avian influenza A viruses are classified as low pathogenicity avian influenza viruses (LPAIVs), or as high pathogenicity avian influenza viruses (HPAIVs) based on their pathogenicity in poultry^1,2^. The major determinant of virus pathogenicity in poultry is the cleavability of the surface glycoprotein haemagglutinin (HA). The HA of LPAIVs has a single basic amino acid that is cleaved by trypsin-like proteases located in the lung and intestinal tract of birds. By contrast, the HA of HPAIV has a multibasic cleavage site (MBCS) that can be proteolytically processed by ubiquitous furin-like proteases present in all cells of the organism^3,4^, enabling the virus to spread systemically and to cause severe disease^5,6^. Consequently, HPAIVs are a major threat to poultry^7^ and wild birds^8^. The LPAIV-to-HPAIV switch is also associated with an increased zoonotic risk, both for the likelihood of human contact and potential pathogenesis^7,9^.

Natural acquisition of an HA MBCS has only been described for viruses belonging to the H5 and H7 subtypes^1,2^ (with the exception of one H4 virus^10^). This observation strongly suggests that the HA of H5 and H7 viruses is either more prone to acquire mutations in the HA cleavage site and/or that it is more likely to gain a phenotypic advantage upon accumulation of basic amino acids at the HA cleavage site. Introduction of a MBCS via reverse-genetic manipulation in the HA of viruses belonging to other subtypes (H2, H4, H6, H8, H9 and H14) conferred an HPAIV phenotype to these viruses^11–13^. These studies demonstrate that the HAs of other subtypes support a HPAIV phenotype once they have a MBCS and strongly suggest that the restriction of naturally occurring HPAIV to H5 and H7 subtypes could be due to a predisposition of H5 and H7 HA subtypes to accumulate nucleotide mutations in the sequence encoding the HA cleavage site.

The acquisition of an HA MBCS proceeds either via viral RNA-dependent RNA polymerase (RdRp)-mediated errors causing nucleotide substitutions and/or insertions, or via non-homologous recombination that adds several codons originating from either viral or host RNAs^1,2^. In this study, we explored the mechanism underlying MBCS acquisition via nucleotide insertions, which account for 9 out of 15 reported H5 HPAIV emergence events and 22 out of 42 reported H7 HPAIV emergence events. Previous work proposed that insertions could be due to the formation of stationary secondary stem-loop RNA structures^14–17^ or of transient secondary RNA structures forming outside of the viral RdRp via the interaction between the 3′ and 5′ regions of the template RNA strand^18^, as these structures could impede viral RdRp processivity and promote nucleotide duplications causing insertions (Fig. 1a). By combining experimental evolution, analyses of natural emergence events and modelling methods, we exclude the contribution of these predicted secondary RNA structures as determinant drivers of HPAIV emergence. By contrast, we demonstrate that the HA cleavage-site RNA sequence, and more precisely the thermodynamic stability of the product–template duplex formed within the viral RdRp catalytic site, is the principal determinant of nucleotide insertions. Furthermore, we developed a mathematical model that predicts the risk of acquiring an HA furin-dependent MBCS via insertions and thus provide the public- and animal-health communities with an operational tool to identify H5 and H7 sequences with elevated risk of evolving towards HPAIV.

**Fig. 1.**
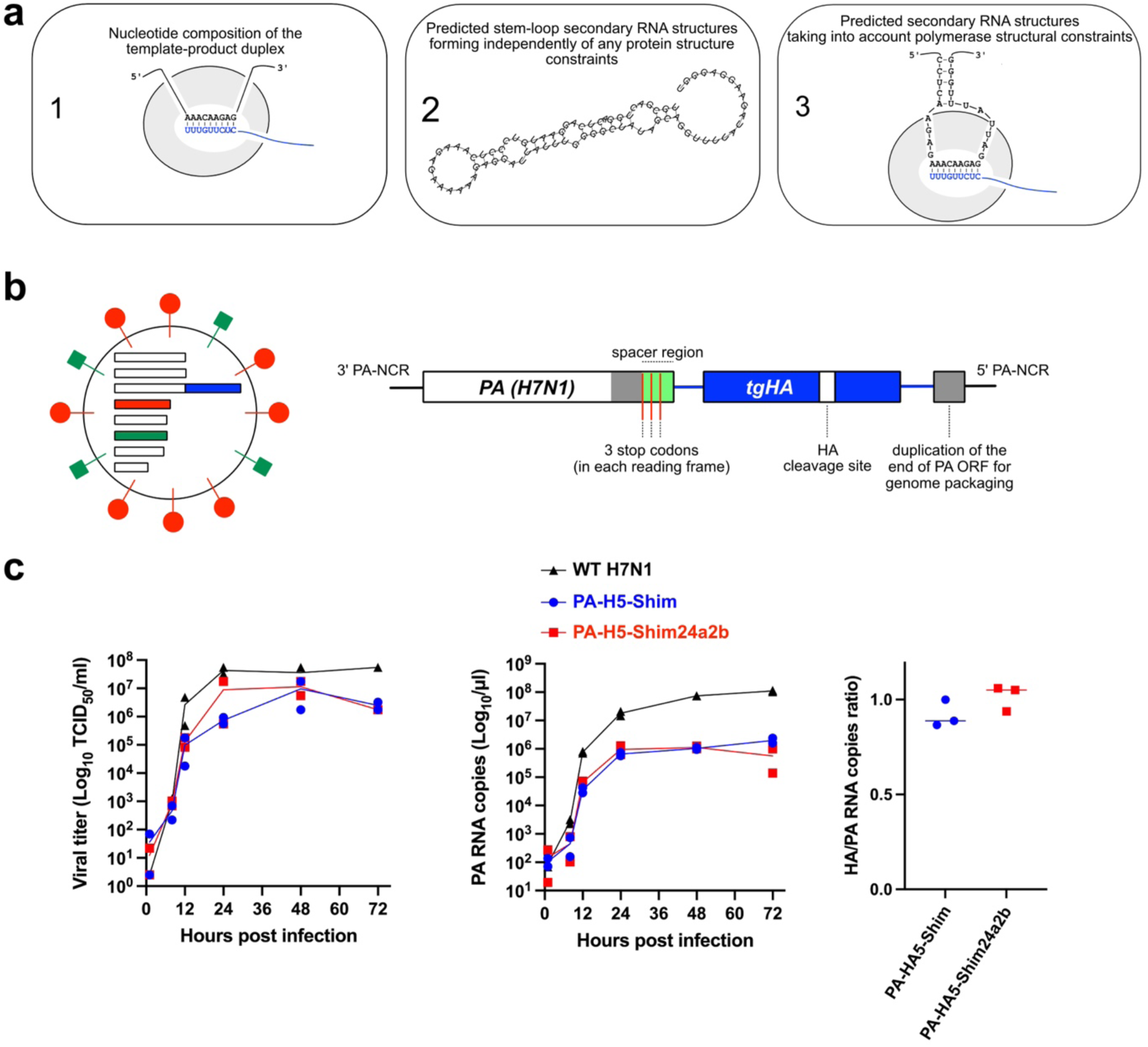
Experimental framework to study HA cleavage-site insertions. **a)** Schematic representation of hypothesized mechanisms underlying nucleotide insertions at the HA cleavage site. Viral RdRp-mediated insertions could be influenced by (1) Local nucleotide composition of the product–template duplex; (2) Formation of stationary secondary stem-loop RNA structures; (3) transient secondary RNA structures forming outside of the viral RdRp through interactions between the 3′ and 5′ ends. **b)** Design of the PA–HA experimental system. Schematics of recombinant H7N1 influenza virus with a PA segment containing the HA of interest inserted after the PA open reading frame as a non-translatable HA (tgHA). **c)** Characterization of the PA-HA system. Growth properties of the wild-type H7N1, PA-H5-Shim and PA-H5-Shim24a2b viruses measured by viral titre (left) and PA RNA copy number (centre), ratio of HA to PA RNA levels, indicating stability of the tgHA (right). Results are presented as means, with individual values from two to three independent experiments.

## Results

### The RNA sequence drives insertion rates

To investigate the impact of specific RNA sequence properties directly on viral RdRp-mediated nucleotide insertions in the HA cleavage site, we developed an original experimental system in which the HA of interest is inserted as a non-translatable transgene after the PA open reading frame using reverse-genetics to produce an infectious H7N1 virus (Fig. 1b). In the PA–HA system, the HA sequence is replicated without direct selection on HA protein function, because a functional HA is supplied by the authentic H7 HA segment. This allows mutations to be examined without directly altering the function of the expressed viral HA. To validate the PA-H5 system, we first produced the PA-H5-Shim and PA-H5-Shim24a2b viruses, containing respectively the H5 of A/whistling swan/Shimane/499/83 (H5N3) parental strain (H5-Shim) and its mutant version (H5-Shim24a2b)^17,19^. These H5 differ by only two nucleotides in the HA cleavage site sequence. H5-Shim24a2b was previously shown to accumulate significantly higher levels of nucleotide insertions than the parental H5-Shim and to evolve via the insertion of a GAA triplet to a high pathogenicity H5 with an elongated MBCS containing the amino acids RRKKR found in several HPAIV^17,19^.

The PA-H5-Shim and PA-H5-Shim24a2b viruses grew to similar levels as the wild-type H7N1 virus. The transgenic H5 was replicated and stable along successive passages (Fig. 1c) and was not detected as a protein (fig. S1). To determine insertion profiles in the transgenic HA, we used Illumina-based single-strand consensus sequencing^20^, which we determined to accurately detect nucleotide insertions from a 2×10^-6^ threshold (fig. S2). In addition, in order to correct for reverse-transcriptase (RT) mediated errors and for errors potentially introduced by the human RNA polymerase I (RNA pol I) during the virus rescue process, for each HA sample, we subtracted nucleotide insertions detected in parallel control experiments performed with the corresponding HA transcribed from human RNA polymerase I (fig. S2). As this workflow was systematically applied, the results presented throughout this work accurately report on the identity and frequencies of nucleotide insertions caused by the viral RdRp.

We analysed viral RNA collected from the supernatants of canine MDCK cells infected with PA-H5-Shim or PA-H5-Shim24a2b at a multiplicity of infection (MOI) of 10^-3^. For clarity, insertion profiles are shown for a focused 19-nucleotide window spanning positions 1012– 1030 of the H5 HA cleavage-site–encoding sequence (Fig. 2a, top panels). In each panel, the HA cleavage-site nucleotide sequence is shown on the X-axis, with the eight-adenine tract of PA-H5-Shim24a2b highlighted in red. Histogram bars represent the average insertion frequency at each position, calculated from three independent experiments. Inserted sequences are indicated by letters above each position, with their vertical placement reflecting insertion frequency reported on the Y-axis. Only insertions detected above the 2×10^-6^ sequencing detection threshold are shown. For PA-H5-Shim, only rare single-nucleotide insertions were detected above this threshold, with a single A or G inserted at positions 1013, 1018 or 1019 (Fig. 2a, top left). By contrast, PA-H5-Shim24a2b displayed markedly higher insertion frequencies at multiple positions, with the highest frequencies at positions 1013, 1016 and 1017 (Fig. 2a, top right), confirming previous observations made with this H5 sequence in an A/Puerto Rico/8/1934 (H1N1) backbone^17^. In PA-H5-Shim24a2b, we detected a nine-nucleotide long insertion at position 1013. Position 1017 displayed an insertion frequency reaching 10^-1^, predominantly consisting of single-A insertions, followed by AA and AAA insertions (Fig 2a, top right). Position 1016 displayed several guanine- and adenine-containing insertions, notably the GAA triplet insertion previously shown to convert the parental RKKR cleavage site of Shim24a2b into the elongated RRKKR MBCS found in several HPAIVs^17,19^.

**Fig. 2.**
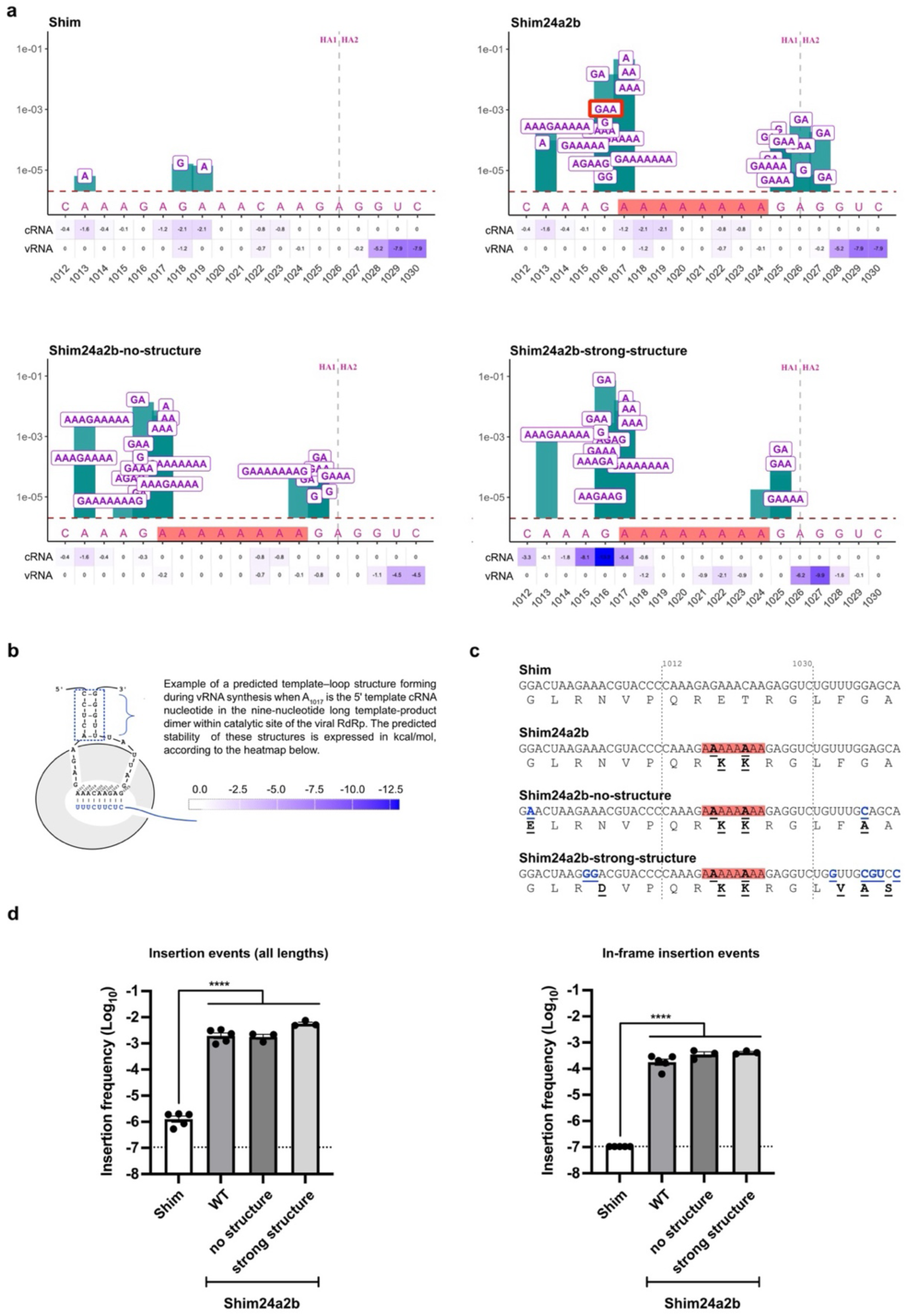
Predicted transient RNA structures are not required for high-frequency HA cleavage-site insertions. **a)** Nucleotide insertion profiles in the HA cleavage-site–encoding sequence of PA-H5-Shim, PA-H5-Shim24a2b, PA-H5-Shim24a2b-no-structure and PA-H5-Shim24a2b-strong-structure. Insertions are shown across a 19-nucleotide window corresponding to positions 1012–1030 of the H5 HA cleavage site. Bars indicate the average insertion frequency at each position from three independent experiments. Letters indicate the nucleotide composition of each insertion, positioned according to its frequency. The horizontal dashed line indicates the sequencing detection threshold of 2×10^-6^ per position, and the vertical dashed line indicates the HA cleavage site. Heatmaps below each X-axis show the predicted stability of template–loop RNA structures during cRNA or vRNA synthesis, with darker blue indicating more stable structures. The GAA triplet insertion generating an elongated MBCS encoding the RRKKR motif found in several H5 HPAIVs is circled in red. The eight-adenine tract found in Shim24a2b and its variants is highlighted in red. **b)** Schematic representation of a predicted template–loop RNA structure forming outside the viral RdRp, together with the heatmap scale used to represent predicted stability in kcal/mol. **c)** Nucleotide and amino acid alignments of the Shim H5 variants. Underlined nucleotides and amino acids indicate substitutions relative to PA-H5-Shim. The eight-adenine tract is highlighted in red. Nucleotides in blue indicate substitutions introduced to alter predicted template–loop stability. **d)** Quantification of insertion frequencies across the HA cleavage-site window. Left, insertion events across all lengths. Right, in-frame insertion events. For each independent experiment, insertion frequencies were averaged across the 19-nucleotide window. Bars show mean ± SEM; points represent independent experiments performed in MDCK and DF-1 cells for PA-H5-Shim and PA-H5-Shim24a2b, and in MDCK cells for PA-H5-Shim24a2b-no-structure and PA-H5-Shim24a2b-strong-structure. Dashed lines indicate the detection threshold of 1.05×10^-7^ after averaging across the 19-nucleotide analysed window. Statistical analyses were performed using one-way ANOVA followed by Tukey’s multiple-comparison test. For clarity, statistical annotations are shown only for comparisons of PA-H5-Shim with PA-H5-Shim24a2b, PA-H5-Shim24a2b-no-structure and PA-H5-Shim24a2b-strong-structure; all three comparisons were significant (**** p<0.0001). All other comparisons were not statistically significant.

The host cell environment had no significant impact, as similar insertion frequencies and patterns were observed in canine MDCK and chicken DF-1 cells (fig. S3). Nor did the viral polymerase background have a detectable effect, as similar results were obtained with the A/WSN/1933 (H1N1), avian H7N1 and H5N8 polymerase complexes (fig. S4). Altogether these results demonstrate that the nucleotide sequence of the HA has a direct and profound effect on viral RdRp-mediated nucleotide insertions in the HA cleavage site.

### Transient RNA structures are not the drivers of insertions

We next investigated potential mechanisms underlying the increased insertion frequency observed in PA-H5-Shim24a2b relative to PA-H5-Shim. Recent studies have proposed that stable transient RNA secondary structures, termed template-loop structures, forming via base pairing between the 3′ and 5′ regions of the template RNA outside the viral RdRp, can impair polymerase processivity^21^ and drive nucleotide insertions at the H5 cleavage site^18^.

To assess this possibility, we applied the sliding-window algorithm described by Funk et al.^18^ to estimate the stability of transient RNA structures that may arise during viral RNA (vRNA) and complementary RNA (cRNA) synthesis in our HAs (Fig. 2b). Predicted template–loop stability profiles are shown as heatmaps beneath each X-axis in Fig. 2a, with darker blue indicating more stable structures. PA-H5-Shim and PA-H5-Shim24a2b exhibited identical predicted stability profiles for both vRNA and cRNA templates. This is consistent with the fact that the two variants differ only at two nucleotide positions that do not contribute to predicted template-loop formation when the cleavage site–encoding region resides within the RdRp catalytic site (Fig. 2c). Thus, the difference in insertion rates between PA-H5-Shim and PA-H5-Shim24a2b cannot be explained by differences in predicted transient structure stability.

Direct mutagenesis confirmed this conclusion. A PA-H5-Shim24a2b-no-structure mutant predicted to lack stable transient structures and a PA-H5-Shim24a2b-strong-structure mutant predicted to form a highly stable template-loop outside the viral RdRp (-13.2 kcal mol^-1^) both retained insertion profiles similar to PA-H5-Shim24a2b, including recurrent GAA insertions that convert the parental RKKR cleavage site into the high-pathogenicity RRKKR motif (Fig. 2a,d). Although PA-H5-Shim24a2b-strong-structure showed an approximately threefold increase in insertion frequency across all insertion lengths relative to PA-H5-Shim24a2b, this effect was minor compared with the ∼600-fold increase observed between PA-H5-Shim24a2b and PA-H5-Shim (Fig. 2d left). Similar trends were observed for in-frame insertion events, which preserve the HA open reading frame, with comparable frequencies across PA-H5-Shim24a2b and its mutant derivatives (Fig. 2d right). No difference in insertion frequency or profile was observed between PA-H5-Shim24a2b and the no-structure mutant.

Together, these results indicate that predicted transient RNA structures forming outside the viral RdRp have, at most, a modest effect on insertion frequency and are not required to generate frequent and reproducible guanine–adenine insertions that create furin-cleavable MBCS.

### Stationary stem-loop structures are not the drivers of insertions

In addition to two substitutions in the H5 CS encoding region forming an eight-nucleotide A-stretch, PA-H5-Shim24a2b exhibited a larger predicted stationary stem-loop RNA structure than PA-H5-Shim presented as insets in Fig. 3a. As such structures were previously linked to insertions in the HA cleavage site^14–17^, we engineered four additional viruses with either the PA-H5-Shim or PA-H5-Shim24a2b stem–loop structure but varying adenine stretch length (Fig. 3b). PA-H5-ShimA1 (Fig. 3a, middle left panel) and PA-H5-ShimA3 (Fig. 3a, bottom left) shared the larger stem–loop of PA-H5-Shim24a2b, whereas PA-H5-ShimA2 (Fig. 3a, middle right panel) and PA-H5-ShimA4 (Fig. 3a, bottom right) retained the smaller structure of PA-H5-Shim. For clarity, the insertion profiles of PA-H5-Shim (Fig. 3, top left) and PA-H5-Shim24a2b (Fig. 3, top right) are reproduced. PA-H5-ShimA1 and A2 showed low insertion frequencies similar to PA-H5-Shim, whereas A3 and A4 exhibited high insertion frequencies comparable to PA-H5-Shim24a2b. Notably, PA-H5-ShimA4 had a small stem– loop yet high insertion frequency, whereas PA-H5-ShimA1 had a large stem–loop but low frequency. These findings indicate that a large stem-loop RNA structure is neither required nor sufficient for frequent insertions at the HA cleavage site.

**Fig. 3.**
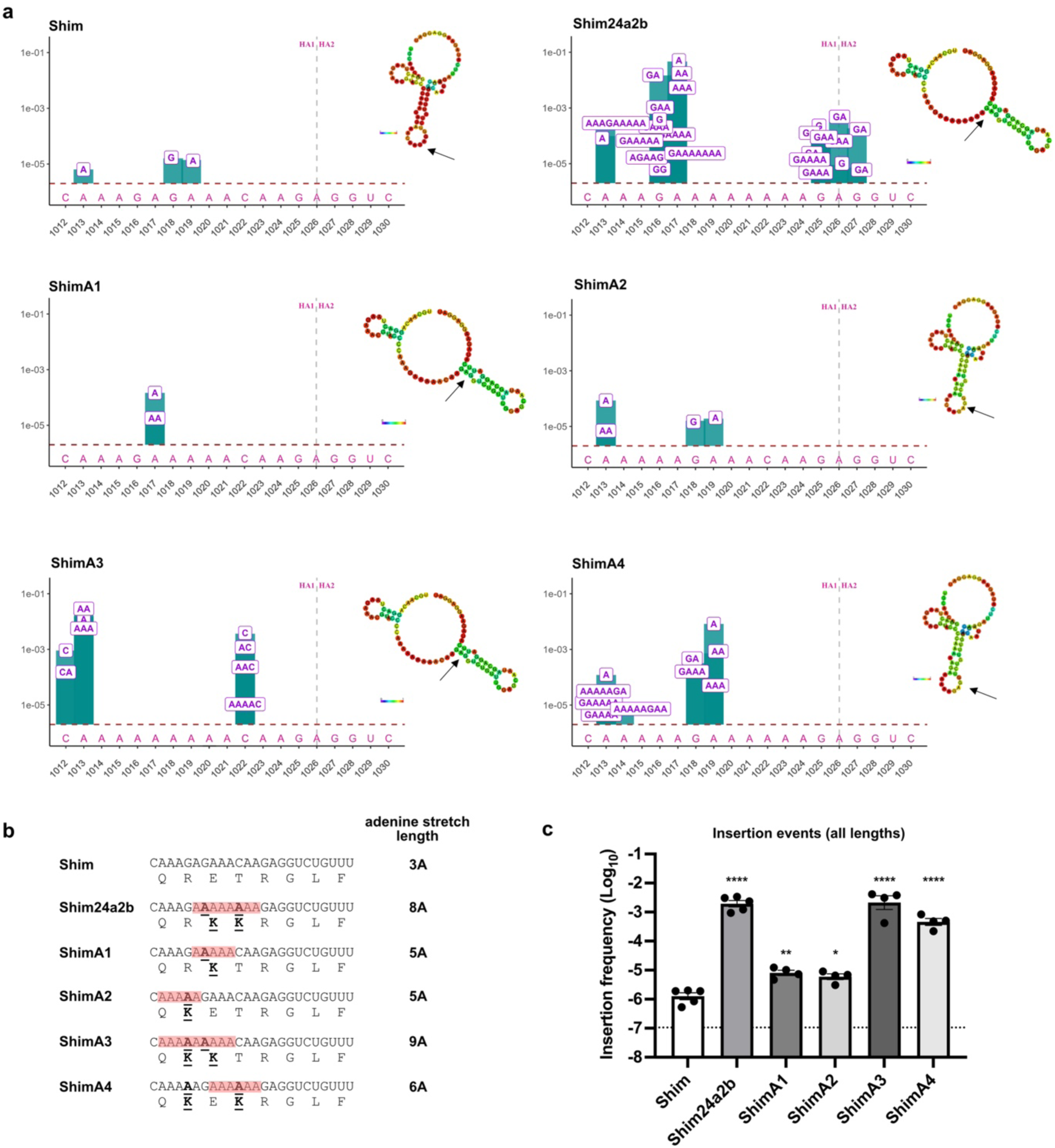
Insertions are driven by the primary RNA sequence independently of stationary stem-loop structures. a) Nucleotide insertion profiles in the HA cleavage site encoding sequence of PA-H5-Shim (top left), PA-H5-Shim24a2b (top right), PA-H5-ShimA1 (middle left), PA-H5-ShimA2 (middle right), PA-H5-ShimA3 (bottom left) and PA-H5-ShimA4 (bottom right). Insertions in a 19-nucleotide window corresponding to nucleotides 1012 to 1030 of the H5 cleavage site are represented in the graphs. The histogram bars represent the average frequency of insertions at each position calculated from three to five independent experiments. The dotted horizontal line corresponds to the 2 × 10⁻⁶ sequencing detection threshold. The vertical dotted line indicates the site of HA cleavage. Predicted stationary stem–loop RNA structures for each H5 sequence are visualized using the interactive minimum free energy drawing provided by ViennaRNA RNAfold server and displayed as insets in each graph with colours assigned by base pair probabilities, with red indicating the highest base-pairing probability and blue the lowest. b) Nucleotide and protein sequence alignments of the different ShimH5 variants. Underlined nucleotides and amino acids indicate substitutions compared to the reference Shim sequence. The adenine stretches are highlighted in red. c) Quantification of insertion events across all lengths within the HA cleavage-site window. For each independent experiment, insertion frequencies were averaged across the 19-nucleotide window. Bars show mean ± s.e.m.; points represent independent experiments. Dashed lines indicate the detection threshold of 1.05×10-7 after averaging across the 19-nucleotide analysed window. Statistical analyses were performed using one-way ANOVA followed by Tukey’s multiple-comparison test. Statistical annotations indicate comparisons of the different variants with PA-H5-Shim; *: P <0.05, **: P <0.01, ****: P <0.0001.

### The primary RNA sequence is the main driver of insertions

By contrast, H5 HA sequences with high insertion frequencies consistently contained a stretch of at least six consecutive adenines within the cleavage site-encoding region - six in ShimA4, eight in Shim24a2b and nine in ShimA3 (Fig. 3b,c) - implicating the primary nucleotide sequence, and particularly the presence of a tract of at least six consecutive adenines, as the dominant determinant of insertion.

To test whether an adenine tract can promote insertions outside the H5 sequence context, we generated a PA-HA virus carrying the HA of A/Puerto Rico/8/1934 (H1N1) (PR8). The wild-type H1 sequence showed only rare single-adenine insertions at position 1013, whereas introduction of a six-adenine stretch increased insertion frequency by approximately 100-fold, with insertions predominantly comprising one to three adenines (**fig. S5**). These results demonstrate that altering the primary sequence alone is sufficient to markedly enhance nucleotide insertion.

### Product–template duplex thermodynamics predicts polymerase slippage

Analysis of insertion diversity revealed that the inserted nucleotide sequences invariably corresponded to duplications of the RNA template, consistent with polymerase slippage as the underlying mechanism^14–16^. Polymerase slippage has been well described as a mechanism leading to viral transcript editing in *Potyviridae*^22,23^, *Filoviridae*^24–26^ and *Paramyxoviridae*^27,28^. This process involves dissociation of the original product–template RNA duplex within the catalytic site of the viral RdRp and reannealing of the nascent RNA at a downstream site on the template. Upon reannealing, synthesis resumes, generating an insertion at the 3′ end of the extending RNA product^29,30^. The inserted nucleotides are complementary to the backtracked template region, thereby duplicating it in the product RNA (Fig. 4a).

**Fig. 4.**
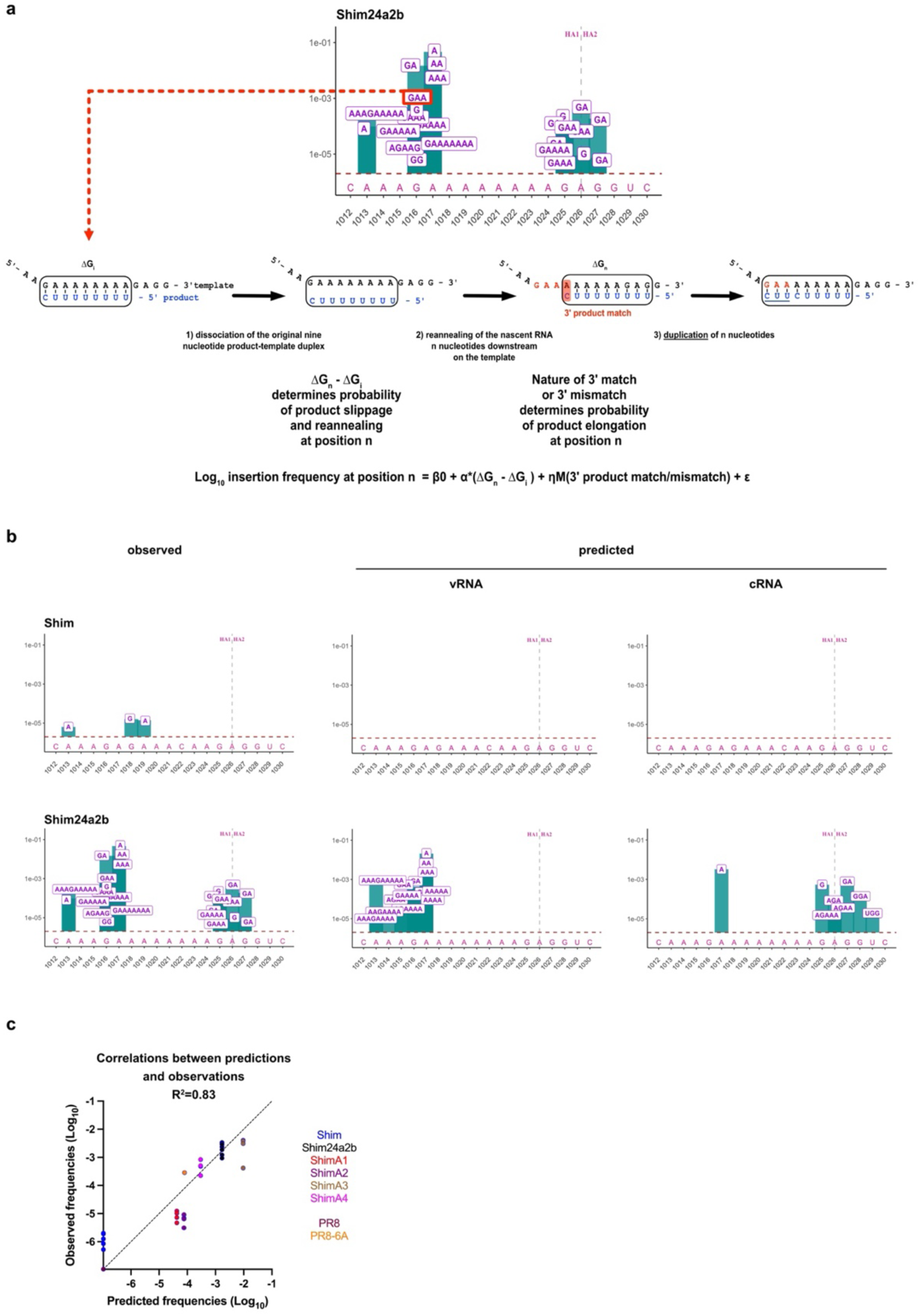
Thermodynamic modelling of RdRp slippage predicts insertion frequency and composition. a) Thermodynamic model of polymerase slippage underlying nucleotide insertions. The PA-H5-Shim24a2b insertion profile is used to illustrate the origin of the GAA insertion at position 1016. This insertion results from a duplication of the complementary CUU triplet during vRNA synthesis due to RdRp slippage, which proceeds according to the following thermodynamic principles. The original nine-nucleotide product–template duplex formed between the cRNA template (in black) and the nascent vRNA product (in blue) has a thermodynamic stability ΔGi. Reduced stability of this duplex favours transient dissociation of the product from the template. The nascent RNA product can then realign n nucleotides downstream of its original position, forming a new product–template duplex with thermodynamic stability ΔGn. Higher stability of this realigned duplex promotes reannealing. The difference in thermodynamic stability between the realigned and original duplexes (ΔΔG= ΔGn-ΔGi) is used to estimate the probability of product slippage and reannealing at position n. From this realigned position, the nature of the 3′ terminal match or mismatch influences elongation efficiency. These two parameters are integrated into the regression component of HPAIVpredict to estimate insertion frequencies. b) Comparison of experimentally observed insertion profiles and HPAIVpredict-predicted profiles for PA-H5-Shim and PA-H5-Shim24a2b. Observed profiles are shown on the left, whereas predicted insertion profiles are shown separately for vRNA and cRNA synthesis. The model captures the position, identity and relative frequency of insertion events, including recurrent triplet insertions in PA-H5-Shim24a2b. The horizontal dashed line indicates the sequencing detection threshold, and the vertical dashed line indicates the HA cleavage site. c) Correlation between HPAIVpredict-predicted and experimentally observed average insertion frequencies across all HA sequences included in model training. For each HA sequence, HPAIVpredict-predicted insertion frequencies were averaged across the 19-nucleotide window spanning the HA cleavage-site-encoding sequence to generate a single predicted value, shown on the x-axis. Experimentally observed insertion frequencies were averaged across the same window for each independent experiment, and individual replicate values are shown on the y-axis. Predicted and observed insertion frequencies showed good agreement across several orders of magnitude (Pearson R² = 0.83).

Given the evidence for a primary nucleotide sequence-driven mechanism, we reasoned that RdRp slippage could be modelled using the thermodynamic stability of product–template duplexes formed at successive steps of the replication process. Disruption of the original duplex should be favoured by thermodynamically unstable product–template interactions, whereas reannealing should depend on the stability of alternative duplexes formed after displacement of the nascent product RNA. In addition, prior work has shown that a base-pair mismatch at the 3′ end of the product impairs elongation^31^, suggesting that the presence or absence of a terminal match could influence insertion efficiency from a reannealed duplex.

We first developed a multiple linear regression model incorporating two predictors: (i) the difference in stability between the original and backtracked nine-nucleotide product–template duplexes^32,33^ (ΔΔG), and (ii) the nature of the 3′ terminal match or mismatch in the reannealed duplex^34–36^ (Fig. 4a). Trained on insertions experimentally observed in the H5-Shim series and H1-PR8 variants (Figs. 2, 3 and S5), the model explained 69% of the variance in insertion frequency (R^2^ = 0.69; F_2,12_ = 13.28; P = 0.0009).

Because slippage to distal positions requires progression through intermediate realigned states, we further considered the thermodynamic accessibility of the full slippage pathway. Predicted insertions whose final backtracked duplex displayed favourable ΔΔG values but whose trajectory required crossing of one or more unstable intermediate duplexes were considered inaccessible, as such intermediates would be expected to act as thermodynamic barriers to progression (fig. S6). Application of this pathway-accessibility constraint increased model specificity by excluding slippage products that were predicted by the regression model but not observed experimentally, including nine-nucleotide insertions (fig. S7).

We refer to the resulting framework, combining two-predictor regression with thermodynamic pathway accessibility, as *HPAIVpredict* (Fig. 4b). Application of *HPAIVpredict* to HA sequences from the H5-Shim and PR8 variants recapitulated the observed insertion profiles and approximate frequencies, as illustrated for PA-H5-Shim and PA-H5-Shim24a2b (fig. S7). *HPAIVpredict* predictions closely matched experimental observations, with good agreement between predicted and observed window-averaged insertion frequencies across several orders of magnitude (R^2^ = 0.83; Fig. 4c), supporting the robustness of the model.

### Decoding natural H5 HPAIV emergence events with *HPAIVpredict*

We next examined all documented natural H5 HPAIV emergence events involving nucleotide insertions to elucidate the mechanisms underlying their acquisition of a MBCS (table S1). Of note, no clear association was detected between HPAIV emergence and the presence of stable predicted transient RNA structures, as only a subset (5/9) of inferred precursor viruses exhibited predicted transient RNA structure stabilities ≤ -7.9 kcal mol⁻¹ (table S1). By contrast, all H5 HPAIVs that emerged via insertions harboured a stretch of five to eight consecutive adenines within the HA cleavage site-encoding sequence (table S1), a feature typically absent from LPAIVs (fig. S8).

To test this hypothesis, we analysed three well-characterized emergence cases (1966 H5N9^37^, 1994 H5N2^14^ and 2006 H5N2^38^) for which LPAIV precursors were identified through co-isolation with HPAIV from the same or neighbouring farms^39^ (highlighted in grey in table S1). For each, we reconstructed the parental LPAIV sequence (RETR cleavage site) and engineered 6A-REKR and 8A-RKKR variants. Insertion profiles were assessed both *in silico* using *HPAIVpredict* and experimentally in the PA-HA system (Fig. 5a and Figs. S9, S10).

**Fig. 5.**
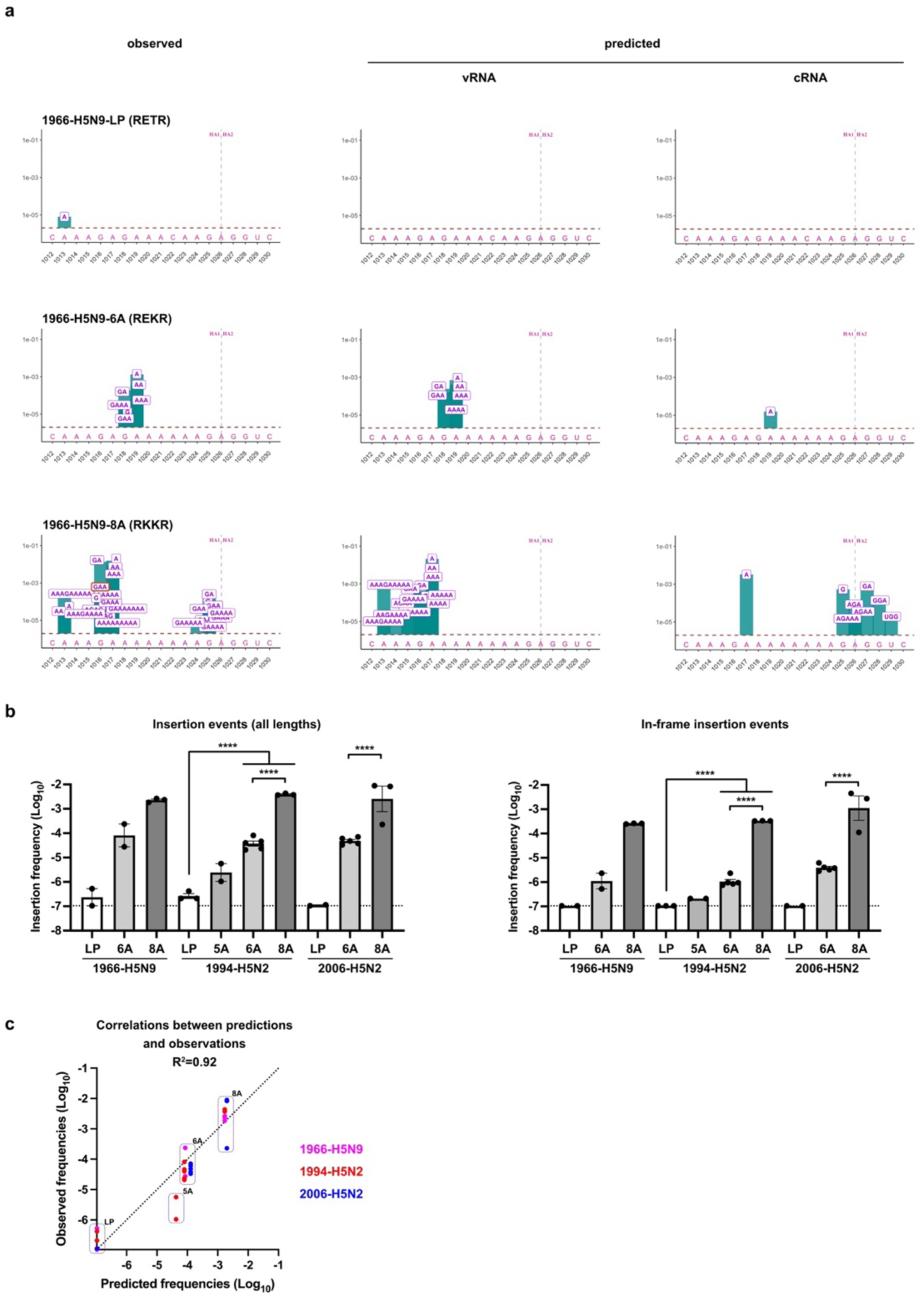
Identification of adenine-rich intermediates underlying insertion-driven H5 HPAIV emergence. a) Comparison of experimentally observed insertions (left) and insertions predicted to occur during vRNA and cRNA synthesis (centre and right, respectively) for the 1966 H5N9 emergence event. The canonical LPAIV precursor (RETR cleavage site), a six-adenine variant (6A-REKR) and an eight-adenine variant (8A-RKKR) were analysed. The histogram bars represent the average frequency of insertions at each position calculated from two to three independent experiments. The letters indicate the nucleotide composition of each type of insertion at the indicated position in the H5 cleavage site sequence, while their position along the y-axis corresponds to the frequency of each type of insertion. The dotted horizontal line corresponds to the 2 × 10⁻⁶ sequencing detection threshold. The vertical dotted line indicates the site of HA cleavage. The GAA triplet insertion generating an elongated MBCS encoding the RRKKR motif found in several H5 HPAIVs is circled in red. b) Quantification of insertion frequencies in the HA cleavage site across the 1966 H5N9, 1994 H5N2 and 2006 H5N2 variants. Left, insertion events across all lengths. Right, in-frame insertion events. For each independent experiment, insertion frequencies were averaged across the 19-nucleotide window. Bars show mean ± s.e.m.; points represent independent experiments. Dashed lines indicate the detection threshold of 1.05×10-7 after averaging across the 19-nucleotide analysed window. Statistical analyses were performed using one-way ANOVA followed by Tukey’s multiple-comparison test. Statistical annotations indicate ****: P <0.0001. c) Correlation between HPAIVpredict-predicted and experimentally observed insertion frequencies across the 1966 H5N9, 1994 H5N2 and 2006 H5N2 variants. For each HA sequence, HPAIVpredict-predicted insertion frequencies were averaged across the 19-nucleotide HA cleavage-site window to generate a single predicted value, shown on the x-axis. Experimentally observed insertion frequencies were averaged across the same window for each independent PA–HA experiment, and individual replicate values are shown on the y-axis. Predicted and observed frequencies show strong agreement (Pearson R² = 0.92).

*HPAIVpredict* did not predict insertions in canonical H5 LPAIV sequences encoding a RETR cleavage site, in agreement with experimental data showing that typical H5 LPAIV cleavage site sequences rarely accumulate insertions (Fig. 5a and fig. S9 and S10). A single C1022A substitution generating a six-adenine stretch (6A-REKR cleavage site) was predicted to substantially increase insertion frequency in all sequences tested, which was confirmed experimentally. Insertions were predicted to occur predominantly during vRNA synthesis. Although triplet insertions were consistently detected in the 6A variants, the inserted codons did not match those observed in naturally emerging HPAIVs, suggesting that such variants are unlikely to represent direct progenitors (Fig. 5a). By contrast, combining the C1022A and G1018A substitutions to generate an eight-adenine stretch (8A-RKKR cleavage site) resulted in a marked increase in predicted insertion frequency, with insertions predicted to occur primarily during vRNA synthesis and, to a lesser extent, during cRNA synthesis, including triplet, hexamer and nonamer insertions compatible with maintenance of the reading frame. These predictions were validated experimentally.

Notably, *HPAIVpredict* predicted the insertion of a GAA triplet at position 1016, which was systematically observed in all 8A-RKKR variants (Fig. 5a and fig. S9 and S10). This insertion converts the 8A-RKKR variants of the 1966 H5N9 and 2006 H5N2 viruses precisely into the HA cleavage site sequences of their natural HPAIV descendants, generating an elongated RRKKR cleavage site. These results provide strong support for the emergence of the 1966 H5N9 CY and 2006 H5N2 EF HPAIVs via an eight-adenine intermediate (Fig. 5a and fig. S9 and S10 and table S1).

These findings identify a stretch of eight consecutive adenines in a RKKR-8A configuration as a major risk factor for H5 HPAIV emergence via insertions. However, shorter adenine tracts (5A-6A), which also strongly increase insertion frequencies (Fig. 5b), may enable MBCS acquisition through less direct or multi-step pathways^14,40^. Extending these observations to natural emergence events not included in model training, *HPAIVpredict* predictions remained in close agreement with experimental measurements (R² = 0.92; Fig. 5c), indicating that the model retains strong predictive performance across diverse sequence contexts.

### Decoding natural H7 HPAIV emergence events with *HPAIVpredict*

We next examined documented H7 HPAIV emergence events involving nucleotide insertions to assess whether similar principles apply (table S2). Consistent with observations in H5, only one inferred precursor exhibited a predicted transient RNA structure stability ≤ −3.9 kcal mol⁻¹, further arguing against a determinant role for such structures in HPAIV emergence (table S2). All insertion-driven H7 emergence cases originated from the African-Eurasian-Oceania (AEO)-lineage (table S2). The canonical H7 LPAIV sequence in this lineage contains a guanine triplet flanked by adenines^41^, suggesting that G→A substitutions could generate insertion-prone adenine tracts (table S2 and fig. S11). To test this hypothesis, we investigated the mechanisms underlying acquisition of a MBCS of the 1979 H7N7^42^ and the 2008 H7N7^43^, two independent H7 HPAIVs that emerged via insertions that unequivocally correspond to template duplications. Because H7 sequences could not be rescued in the PA-HA system, insertion profiles were assessed using a minigenome assay alongside *HPAIVpredict* predictions.

To investigate the 2008 H7N7 HPAIV emergence, we generated the parental LPAIV precursor carrying a canonical H7 cleavage site sequence (LP-PKGR)^43^. *HPAIVpredict* did not predict insertions in this background, in close agreement with experimental observations (Fig. 6a). Introduction of G994A and G995A substitutions to generate a 4A-PKKR variant was predicted to moderately increase insertion frequency, including insertion of an AAA triplet, which was experimentally detected at very low frequency. Further extension to a 5A-PKRR variant (G993A and G994A substitutions) was predicted to increase insertion frequency, notably including insertion of a GAA triplet at position 998, which was experimentally detected at low frequency. Notably, this insertion generates an elongated PKRRR MBCS previously reported in AEO-derived HPAIVs^44^, suggesting that such variants may represent direct evolutionary intermediates. Replacement of the guanine triplet with an adenine triplet to generate an 8A-PKKR variant was predicted to cause a marked increase in insertion frequency, with insertions occurring during both vRNA and cRNA synthesis. These predictions were confirmed experimentally, although insertion frequencies were slightly lower than predicted.

**Fig. 6.**
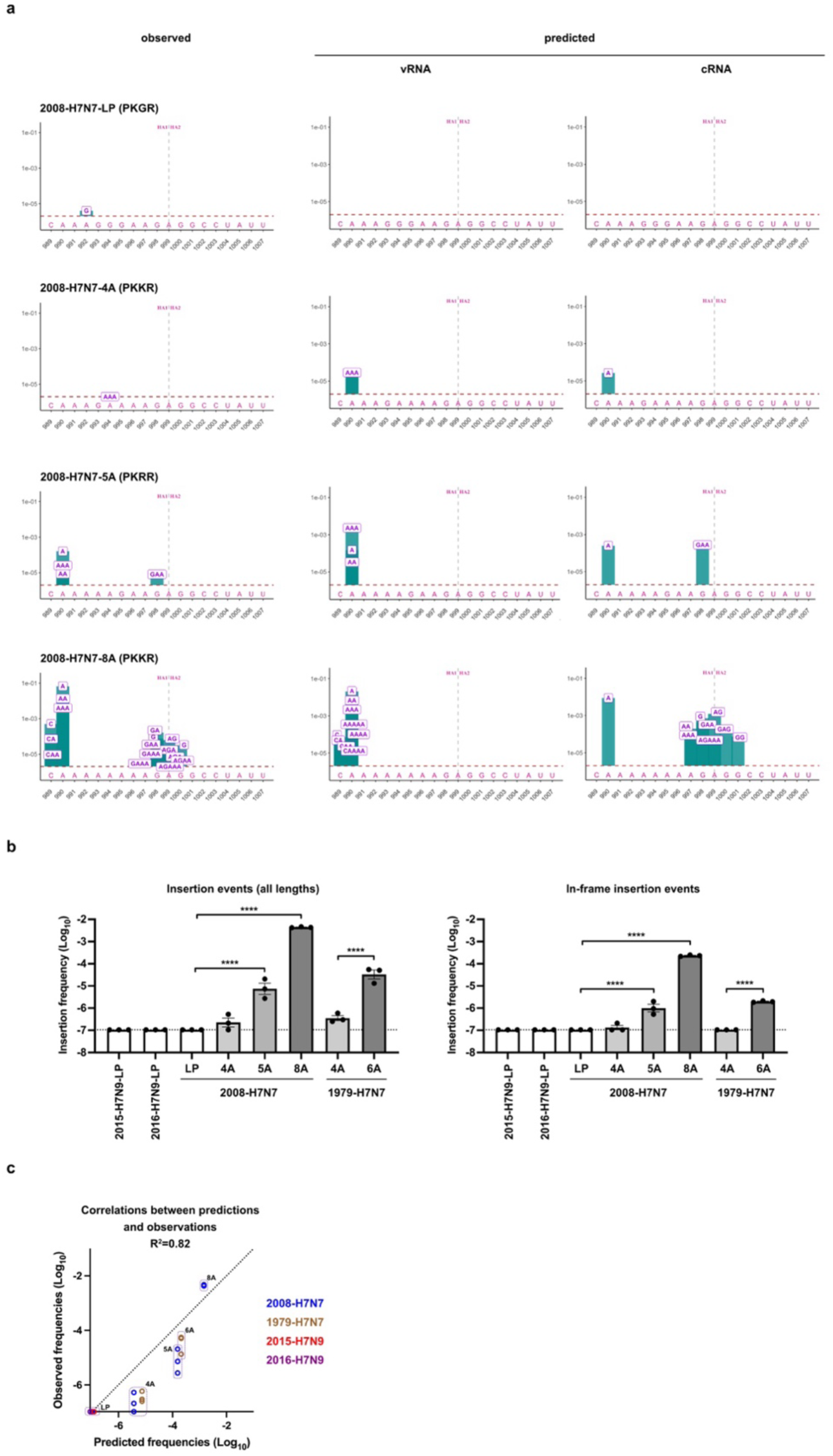
Identification of adenine-rich intermediates underlying insertion-driven H7 HPAIV emergence. a) Comparison of experimentally observed insertions (left) and insertions predicted to occur during vRNA and cRNA synthesis (centre and right, respectively) for the 2008 H7N7 HPAIV emergence event. The canonical LPAIV precursor (PKGR cleavage site), a four-adenine variant (4A-PKKR), a five-adenine variant (5A-PKRR) and an eight-adenine variant (8A-PKKR) were analysed. The histogram bars represent the average frequency of insertions at each position calculated from two to three independent experiments. The letters indicate the nucleotide composition of each type of insertion at the indicated position in the H7 cleavage site sequence, while their position along the y-axis corresponds to the frequency of each type of insertion. The dotted horizontal line corresponds to the 2 × 10⁻⁶ sequencing detection threshold. The vertical dotted line indicates the site of HA cleavage. b) Quantification of insertion frequencies in the HA cleavage site across the 2008 H7N7 and 1979 H7N7 variants, and the 2015 H7N9 and 2016 H7N9 LPAIV. Left, insertion events across all lengths. Right, in-frame insertion events. For each independent experiment, insertion frequencies were averaged across the 19-nucleotide window. Bars show mean ± s.e.m.; points represent independent experiments. Dashed lines indicate the detection threshold of 1.05×10-7 after averaging across the 19-nucleotide analysed window. Statistical analyses were performed using one-way ANOVA followed by Tukey’s multiple-comparison test. Statistical annotations indicate ****: P <0.0001. c) Correlation between HPAIVpredict-predicted and observed insertion frequencies across the 2008 H7N7 and 1979 H7N7 variants, and the 2015 H7N9 and 2016 H7N9 LPAIV. For each HA sequence, HPAIVpredict-predicted insertion frequencies were averaged across the 19-nucleotide HA cleavage-site window to generate a single predicted value, shown on the x-axis. Experimentally observed insertion frequencies were averaged across the same window for each independent minigenome experiment, and individual replicate values are shown on the y-axis. Predicted and observed frequencies show good agreement (Pearson R² = 0.82).

In the case of the 1979 H7N7 HPAIV emergence, sequence comparisons indicated that the HPAIV HA was closely related to an LPAIV bearing an atypical cleavage site sequence containing a four-adenine stretch. We therefore generated this putative precursor (4A-PKGR) and a mutant carrying additional guanine-to-adenine substitutions to produce a six-adenine stretch (6A-PKKR). *HPAIVpredict* predicted only rare single-adenine insertions in the 4A-PKGR context, in agreement with experimental observations (fig. S12). Extension of the adenine tract to generate the 6A-PKKR variant was predicted to increase insertion frequency, including insertion of an AAA triplet, which was experimentally detected at low frequency. As observed for the 2008 H7N7 context, insertions were predicted to occur during both vRNA and cRNA synthesis. Finally, *HPAIVpredict* specificity was further evaluated using two additional H7 LPAIV sequence contexts (2015 H7N9 and 2016 H7N9), for which no insertions were predicted. Experimentally, no insertions were detected in the 2016 H7N9 context, whereas a single rare C insertion was detected in the 2015 H7N9 context (fig. S13).

Across H7 sequence contexts, insertion frequency increased progressively with adenine-tract length (Fig. 6b). *HPAIVpredict* captured the majority of insertion patterns, although predicted frequencies tended to be higher than those observed experimentally (R² = 0.82; Fig. 6c), possibly reflecting differences between the minigenome system used for H7 and the PA-HA system used for H5.

Together, these analyses reveal key distinctions between insertion-prone H5 and H7 sequences. First, whereas H5 sequences typically require at least a six-adenine tract to support nucleotide triplet insertions, H7 sequences can do so with five adenines. Second, while insertions predominantly occur during vRNA synthesis in H5, they occur at comparable levels during vRNA and cRNA synthesis in H7, consistent with sequence features that facilitate reannealing in both directions. These findings highlight the dual sequence constraints governing insertion: an adenine-rich tract promoting duplex destabilization, and a surrounding nucleotide context enabling reannealing of the slipped product.

To test whether these sequence requirements could confer insertion susceptibility outside H5 and H7, we analysed a prototypic H12 virus, a subtype phylogenetically distant from H5 and H7 for which elongated HA cleavage sites have not been reported. The wild-type H12 HA showed only rare insertions in the minigenome assay (fig. S14). By contrast, introduction of five substitutions that generated an adenine-rich tract with flanking nucleotides that *HPAIVpredict* predicted to favour reannealing rendered the H12 sequence insertion-prone (fig. S14). These results further support the conclusion that insertion susceptibility requires both an adenine-rich tract and a surrounding nucleotide context permissive to polymerase slippage.

### Database-scale prediction identifies sequence backgrounds at risk

Finally, we applied *HPAIVpredict* to all GISAID LPAIV H5 and H7 HA sequences available in our dataset, collapsing identical 44-nucleotide cleavage-site windows into 344 H5 and 286 H7 representative sequence contexts. For each context, we predicted insertion frequencies during vRNA and cRNA synthesis, reconstructed in-frame insertion products and scored furin-cleavability with ProP^45^. HA cleavage site sequences with ProP furin-cleavability scores equal to or greater than the minimum score observed among naturally occurring H5 or H7 HPAIV cleavage sites were classified as predicted furin-cleavable MBCS. The complete H5 and H7 prediction datasets are provided as supplementary workbooks (data S1 and data S2). The “WT” sheets contain the H5 and H7 sequences retrieved from GISAID.

Most H5 LPAIV sequence backgrounds exhibited three- to four-adenine tracts and were not predicted to generate insertions above the detection threshold (data S1). Rare LPAIV sequence backgrounds harbouring 5A tracts, often associated with KETR- or RKTR-like cleavage site motifs, were predicted to accumulate low-frequency single-adenine insertions, consistent with experimental observations. Rare sequence backgrounds containing a 6A tract and a REKR cleavage site displayed substantially higher predicted insertion frequencies, although none of the resulting in-frame insertions generated a predicted furin-cleavable MBCS. By contrast, a small number of RKKR sequence backgrounds containing a 7A tract, together with a single RKKR sequence background containing an 8A tract (A/chicken/Jalisco/CPA-04948-23/2023), were predicted to generate one or more in-frame insertions yielding predicted furin-cleavable MBCS.

Most H7 LPAIV sequence backgrounds exhibited two- to four-adenine tracts and were not predicted to generate insertions above the detection threshold (data S2). Notable exceptions included rare 4A-PKKR sequence backgrounds predicted to accumulate adenine-triplet insertions generating predicted furin-cleavable MBCS, consistent with our experimental data. Similarly, rare 5A-PKRR sequence backgrounds were predicted to favour GAA triplet insertions producing predicted furin-cleavable MBCS. Importantly, Australian H7 sequence backgrounds (including A/duck/Tasmania/277/2007 and A/duck/Victoria/512/2007) carried a unique 5A tract associated with a PRKR cleavage site and were predicted to acquire GAA triplet insertions leading to predicted furin-cleavable MBCS. Additional rare H7 sequence backgrounds carrying 5A tracts, as well as unique 6A- and 7A-containing backgrounds, were predicted to accumulate adenine-triplet insertions generating furin-cleavable sites, again consistent with the experimental observation that specific H7 nucleotide contexts can support MBCS-generating insertions.

Analyses of natural HPAIV emergence events and our experimental data suggest that insertion-driven transitions to HPAIV are preceded by substitutions to adenine that generate RNA sequences more permissive to viral RdRp slippage. To model this process, we simulated substitutions to adenine individually and in combination across all H5 and H7 sequence backgrounds. The positions of simulated adenine substitutions and their predicted consequences for insertion frequency and furin-cleavable MBCS acquisition are reported in dedicated sheets of the H5 and H7 supplementary workbooks.

Most canonical H5 and H7 sequence backgrounds were predicted to reach insertion-prone configurations capable of generating a furin-cleavable MBCS after substitution to adenine at specific positions, or combinations thereof. However, some H5 sequence backgrounds reached RKKR cleavage sites with either a 4A tract (for example, A/chicken/Coahuila/IA20/2011) or a 7A tract (for example, A/chicken/Jalisco/18-13/2013) but were not predicted to accumulate in-frame insertions leading to a furin-cleavable MBCS. Notably, most Australian H5 sequence backgrounds failed to reach a configuration prone to acquire in-frame insertions generating a MBCS, even after combinations of three adenine substitutions (for example, A/duck/Werribee/2514/2007), consistent with the absence of confirmed H5 high-pathogenicity emergence events in Australia^2,46^. Similarly, most American-lineage H7 sequence backgrounds failed to reach insertion-prone configurations even after simulated substitutions generating extended adenine tracts, such as the 6A variant of A/chicken/New_York/1995. This is consistent with the observation that American H7 HPAIVs predominantly emerge through non-homologous recombination rather than nucleotide insertions^47,48^.

Collectively, these analyses indicate that adenine-tract length alone does not determine insertion risk. Instead, insertion susceptibility and functional MBCS potential depend on the combined effects of adenine-tract length, local nucleotide context permissive to polymerase slippage and reannealing, and the capacity of resulting in-frame insertions to encode basic amino acids.

## Discussion

The restriction of HPAIV emergence to H5 and H7 subtypes and the mechanisms underlying HA MBCS acquisition have remained unresolved for decades^2^. Previous studies proposed that stationary stem-loop RNA structures^14–17^ or transient secondary RNA structures forming outside the viral RdRp through interactions between the 3′ and 5′ regions of the template RNA strand^18^ may promote acquisition of an HA MBCS via nucleotide insertions by impairing viral RdRp processivity. Here, combining experimental evolution with bioinformatic analyses, we rule out a determinant role for predicted secondary RNA structures. This conclusion is also supported by analyses of natural H5 and H7 HPAIV emergence events and their precursors, which do not consistently exhibit predicted transient RNA structures with stabilities compatible with RdRp trapping (5/9 for H5 and none for H7). Instead, we provide convergent evidence that insertion risk at the HA cleavage site is primarily dictated by the local nucleotide sequence within the viral RdRp catalytic site during replication.

We show that the thermodynamic stability of the product–template duplex, together with the surrounding nucleotide context, governs insertion events at the HA cleavage site. Nucleotide insertions arise through polymerase slippage, during which transient dissociation of the duplex allows the nascent strand to reanneal in a misaligned configuration, generating duplications. Such behaviour is reminiscent of transcriptional editing described in *Potyviridae*, *Filoviridae* and *Paramyxoviridae*^22–29^, where slippage has also been linked to thermodynamically unstable product–template interactions^49,50^. However, whereas transcriptional editing increases mRNA diversity, the insertions described here are heritable, linking intrinsic sequence properties to evolutionary potential.

Our analyses indicate that predicted transient RNA structures forming outside the viral RdRp have, at most, a modest influence on insertion frequency and are not the main drivers of insertion events, in contrast to previous proposals^18^. This discrepancy may partly reflect differences in dynamic range and data visualization: whereas previous analyses used a higher effective detection threshold and linear-scale representation, our ultra-deep sequencing quantified insertion frequencies across several orders of magnitude, represented on a log_10_ scale. When our data are visualized on a linear scale or censored to emulate a higher detection threshold, lower-frequency insertion events become largely invisible and modest differences in the higher-frequency range dominate the representation (**fig. S**15).

Nucleotide insertions at the HA cleavage site differ mechanistically from influenza mRNA polyadenylation. Polyadenylation is driven by a highly reproducible structural constraint at the template terminus, resulting in reiterative adenine insertion (“stuttering”)^51^. By contrast, insertions at the HA cleavage site, which invariably correspond to duplications of the RNA template and are thus consistent with polymerase slippage, occur at comparatively low frequencies, indicating that the underlying determinants act in a probabilistic rather than deterministic manner. Although our data identify local base-pairing thermodynamics as the principal driver of insertion, additional factors - including polymerase conformational flexibility and stochastic movement along the RNA template - may further modulate slippage efficiency.

Building on these principles, we developed *HPAIVpredict*, an algorithm that estimates insertion frequency and composition from the thermodynamic properties of the HA cleavage site sequence. Using this framework, we identified H5 and H7 viruses with adenine-rich cleavage sites and surrounding nucleotide contexts permissive to polymerase slippage, suggesting that some naturally circulating sequences are genetically predisposed to acquire insertion-derived MBCS. Notably, Australian H7 sequences showed elevated potential for triplet insertions leading to MBCS acquisition. Conversely, *HPAIVpredict* also identified sequence backgrounds that remained refractory to insertion even when adenine tracts were extended. In particular, most American-lineage H7 and Australian H5 sequences failed to reach insertion-permissive contexts after simulated substitutions, consistent with the predominance of recombination-driven emergence in American H7 HPAIV^47,48^ and the absence of reported H5 HPAIV emergence in Australia^2,46^. Together, these findings highlight the potential of *HPAIVpredict* to inform sequence-based surveillance strategies aimed at mitigating HPAIV emergence.

Our findings indicate that adenine-rich tracts in the HA cleavage site are a prerequisite for insertion-driven HPAIV emergence^41^. We further identify distinct G→A or C→A substitutions in H5 and H7 that promote the formation of intermediate variants with elevated potential to acquire an elongated, functional MBCS. Restriction of insertion-driven HPAIV emergence to H5 and H7 subtypes likely reflects the convergence of three requirements: a propensity to generate adenine-rich tracts through a small number of substitutions^41^, a nucleotide context permissive to polymerase slippage, and the capacity of inserted nucleotides to encode basic amino acids that enhance HA cleavability. Only when these requirements coincide can insertions generate viable, selectively advantageous high-pathogenicity variants. While this mechanism helps explain why insertion-driven emergence is restricted to H5 and H7 subtypes, the basis for the similar subtype restriction observed for recombination- and substitution-driven pathways remains unresolved.

Strikingly, H5 and H7 sequences predicted to generate in-frame insertions yielding furin-cleavable MBCS are markedly underrepresented in sequence databases, as shown in database analyses presented in data S1 and data S2, despite encoding dibasic HA cleavage sites that can be compatible with HA function, and in some contexts advantageous for HA cleavage^19^. Their scarcity is therefore unlikely to be explained solely by negative selection on HA protein function. Instead, these viruses may represent transient intermediates, present only briefly because of their elevated propensity to acquire insertions leading to elongated MBCS. Once HPAIV variants emerge and gain a strong selective advantage, their immediate precursors may be rapidly outcompeted and remain largely undetected. Consistent with this model, several studies have captured rare intermediate variants harbouring adenine-rich dibasic cleavage sites during emergence events through intensive sequencing of experimental or epidemiologically linked samples^14,19,43,52^.

Comparisons across cell types and viral polymerase backgrounds further support the conclusion that insertion propensity is encoded primarily by the viral RNA sequence. We observed similar insertion profiles in canine MDCK and chicken DF-1 cells, and comparable results across three influenza polymerase complexes, including in minigenome assays in human 293T cells. Together with recent observations that insertion profiles are similar in mammalian and avian cells^18^, these data argue against a major role for species-specific host factors in the molecular generation of HA cleavage site insertions, and suggest that polymerase-background differences are not the dominant determinant in the systems tested. Thus, species-specific differences in HPAIV emergence are more likely to arise from selection acting on variants after MBCS acquisition^39,53,54^, rather than from host-specific control of polymerase slippage.

Together, our findings establish a primary sequence-driven and thermodynamically governed mechanism for nucleotide insertions at the HA cleavage site, providing a mechanistic framework for understanding the emergence of HPAIV via insertions. More broadly, these results provide a foundation for sequence-informed surveillance strategies aimed at mitigating HPAIV emergence at its source and highlight how intrinsic sequence properties can shape evolutionary trajectories.

## Materials and Methods

### Biosafety and regulatory approvals

All experiments involving recombinant influenza viruses and genetically modified organisms were conducted in approved containment facilities in accordance with institutional biosafety procedures and French regulations governing GMO research. The project was reviewed and approved by the French GMO committee under authorization 8756, 10813, 12637, 12640, 13474 and 13586, which covered the generation and use of the recombinant viruses described in this study. The PA–HA viruses used in this study carried HA sequences as non-translatable transgenes and were handled under the approved GMO protocol and procedures in BSL-3 conditions.

### Design of PA-HA viruses and cloning

PA-HA viruses were generated in the genetic background of A/turkey/Italy/977/1999 (H7N1)^55^. To construct these viruses, HA sequences of interest were inserted immediately downstream of the PA stop codon in the reverse-genetics pHW2000-PA plasmid^56^.

A synthetic DNA fragment containing the terminal 60 nucleotides of the PA coding sequence followed by a linker cloning site (LCS) harbouring two BsmBI restriction sites was synthesized in pUC57 by GenScript (pUC57-PA-LCS). The pHW2000-PA and pUC57-PA-LCS plasmids were digested with PvuII and assembled to generate pHW2000-PA-LCS. In this construct, the terminal 60 nucleotides of the PA open reading frame were duplicated and placed upstream of the LCS. This design preserves the contiguous packaging signals located in both the coding and non-coding regions at the 5′ end of the PA segment, as previously described^57^. To reduce the risk of genetic instability caused by repeated PA sequences, 18 synonymous substitutions were introduced into the terminal region of the PA open reading frame using the In-Fusion HD Cloning Kit (Takara).

HA inserts were cloned into pHW2000-PA-LCS. These included HA sequences derived from A/whistling swan/Shimane/499/83 (H5N3) and its variants (Shim24a2b, Shim24a2b-no-structure, Shim24a2b-strong-structure, ShimA1, ShimA2, ShimA3 and ShimA4), HA from A/Puerto Rico/8/1934 (H1N1) and its engineered 6A variant, and HA sequences from the precursors of the 1966 H5N9, 1994 H5N2 and 2006 H5N2 HPAIVs together with their engineered 5A, 6A and 8A variants where indicated. All HA sequences were synthesized as gene fragments by Genewiz, and all cloned constructs were verified by Sanger sequencing.

### Cells and virus production

MDCK, HEK-293T and DF-1 cells were maintained in Dulbecco’s modified Eagle’s medium (DMEM) supplemented with 10% foetal bovine serum and 1% penicillin-streptomycin. Cells were incubated at 37 °C in 5% CO₂.

All experiments with live recombinant viruses were carried out under BSL-3 conditions in the laboratory of the ENVT, INRAE, IHAP, UMR 1225 in Toulouse, France. Viruses were rescued by reverse genetics as previously described^53,56^. For PA-HA viruses, the pHW2000-PA-HA plasmid was used in place of the wild-type pHW2000-PA plasmid. The remaining seven reverse-genetics plasmids were derived from A/turkey/Italy/977/1999 (H7N1). Co-cultures of MDCK and HEK-293T cells, or DF-1 and HEK-293T cells, were transfected with 500 ng of each of the eight pHW2000 plasmids and maintained in Opti-MEM Reduced Serum Medium supplemented with 1% penicillin-streptomycin. TPCK-treated trypsin was added 20 h after transfection to a final concentration of 1 μg ml⁻¹.

At 72 h post-transfection, supernatants containing rescued viruses were collected and clarified by centrifugation to remove cell debris. Viral stocks were generated by amplification of rescued viruses in MDCK cells for viruses obtained from MDCK-HEK-293T co-cultures, or in DF-1 cells for viruses obtained from DF-1-HEK-293T co-cultures. Cells were maintained in Opti-MEM supplemented with 1% penicillin-streptomycin and 1 μg ml⁻¹ TPCK-treated trypsin for 3 days. Virus titres were determined on the corresponding cell type by TCID_50_ assay and calculated using the Spearman-Kärber method.

For passage experiments, viral stocks were used to infect cells at a multiplicity of infection of 0.001 TCID_50_ per cell in viral growth medium consisting of Opti-MEM supplemented with 1% penicillin-streptomycin and 1 μg ml⁻¹ TPCK-treated trypsin. Infections were maintained for 48-72 h, depending on cytopathic effect and monolayer integrity. When approximately 30% of adherent cells remained, supernatants were collected and clarified. Viral RNA was extracted from each sample using the NucleoSpin RNA Virus kit (Macherey-Nagel). RNA obtained after passage 1 was used to assess transgene integrity and stability and to prepare sequencing libraries.

### PA-HA virus characterization

To compare replication kinetics of wild-type and PA-HA viruses, MDCK cells were infected with wild-type A/turkey/Italy/977/1999 (H7N1) (WT H7N1), PA-H5-Shim or PA-H5-Shim24a2b at a multiplicity of infection of 0.0001 TCID_50_. Infections were maintained for 72 h, and supernatants were collected at the indicated time points for virus titration by TCID_50_ assay.

H5 HA protein expression was assessed by western blot. MDCK cells were infected at a multiplicity of infection of 0.1 TCID_50_ per cell for 24 h with recombinant viruses expressing the H7 HA of A/turkey/Italy/977/1999, the H5 HA of A/whistling swan/Shimane/499/83 or its Shim24a2b mutant as functional proteins in a recombinant HPAIV H5N8 background^53^, or carrying the corresponding H5 sequences as non-translatable transgenes in PA-H5-Shim or PA-H5-Shim24a2b viruses. Cells were collected and lysed in RIPA buffer containing 50 mM Tris-HCl pH 8.0, 150 mM NaCl, 1% Igepal CA-630, 0.5% sodium deoxycholate and 0.1% SDS, supplemented with cOmplete Mini EDTA-free protease inhibitor cocktail (Roche).

Membranes were incubated with primary antibodies diluted in TBS-T containing 3% milk: anti-H5 HA (BEI Resources NR-163, 1:1,000), anti-NP (GeneTex GTX125989, 1:5,000) and anti-β-actin (Santa Cruz sc-47778, 1:2,000). HRP-conjugated secondary antibodies were diluted in TBS-T containing 5% milk: anti-goat IgG for HA detection (Sigma A5420, 1:4,000), anti-rabbit IgG for NP detection (Sigma A0545, 1:2,000) and anti-mouse IgG for β-actin detection (Sigma A9044, 1:2,000). Signal was developed using Clarity Western ECL substrate (Bio-Rad) and detected using a Bio-Rad ChemiDoc imaging system.

Integrity of HA transgenes was assessed by RT-PCR, based on the expected size of the transgene-containing amplicon. Transgene stability was further quantified by RT-qPCR using the ratio of HA-transgene vRNA copies to PA vRNA copies, with primer pairs targeting HA and PA, respectively.

### Minigenome assays and human RNA polymerase I controls

For minigenome assays, HEK-293T cells were cultured and transfected using Lipofectamine LTX with PLUS reagent (Invitrogen), according to the manufacturer’s instructions. Cells were transfected with 500 ng of each pcDNA3.1 plasmid encoding the A/WSN/1933 polymerase subunits PB2, PB1 and PA, 500 ng of pcDNA3.1 encoding A/WSN/1933 NP, and 500 ng of a plasmid expressing the HA vRNA sequence of interest from a human RNA polymerase I promoter. The RNA polymerase I HA expression plasmids were generated by deleting the CMV promoter from pHW2000.

HA sequences were derived from A/chicken/England/1158-11406-1/2008 (H7N7), A/chicken/Leipzig/1979 (H7N7) and A/duck/MW04/Vic/2012 (H12N5), together with engineered variants of each sequence. All HA sequences were synthesized as gene fragments by Genewiz and verified by Sanger sequencing after cloning.

For human RNA polymerase I control samples, cells were transfected with the HA expression plasmid alone. Mutations detected in these control RNAs were used to identify and subtract sequence changes arising independently of viral polymerase activity from those detected in PA-HA virus infections or minigenome assays.

Total RNA was extracted 48 h after transfection using the NucleoSpin RNA kit (Macherey-Nagel), according to the manufacturer’s instructions. Residual DNA was removed using ezDNase (Invitrogen). RNA samples were then processed for quality control and sequencing library preparation.

Illumina-based single-strand consensus sequencing of the HA cleavage-site–encoding region Mutability of the HA cleavage site-encoding region was analysed by high-throughput Illumina sequencing using an adapted single-strand consensus sequencing protocol^20^. RNA samples obtained from minigenome assays, RNA polymerase I control transfections and PA-HA virus infections were processed using the same workflow.

For each sample, 5 × 10^6^ viral RNA molecules, estimated by RT-qPCR analyses, were used for cDNA synthesis with AccuScript High-Fidelity Reverse Transcriptase (Agilent). Libraries were generated using three sequential PCR steps, referred to as PCR0, PCR1 and PCR2.

In PCR0, viral cDNA was pre-amplified using Platinum SuperFi II PCR Master Mix (Thermo Fisher). Each reaction contained 5 μl cDNA, corresponding to a theoretical input of 1.25 × 10^6^ single-stranded DNA molecules, 2.5 μl of each 10 μM gene-specific primer flanking the HA cleavage site region, 25 μl PCR master mix and 15 μl nuclease-free water. Cycling conditions were: 98 °C for 30 s; 21 cycles of 98 °C for 10 s, 56 °C for 10 s and 72 °C for 10 s; 72 °C for 5 min; and hold at 4 °C. A fraction of each PCR product was analysed on a 1.5% agarose gel to verify amplicon size and purity. Remaining PCR products were purified using AMPure XP beads (Beckman Coulter) at a 1:1 sample-to-bead volume ratio and quantified using the Quant-iT PicoGreen dsDNA Assay Kit (Thermo Fisher).

PCR0 amplicons were used as templates for PCR1, which introduced unique molecular identifiers (UMIs) and partial Illumina adapter sequences^20^. PCR1 reactions contained 8 μl PCR0 amplicon diluted to 0.5 ng μl⁻¹ and 1 μl each of 10 μM forward and reverse primers containing a template-specific region, an eight-nucleotide random barcode and a partial Illumina adapter sequence at the 5′ end. Cycling conditions were: 98 °C for 30 s; 8 cycles of 98 °C for 10 s, 56 °C for 10 s and 72 °C for 10 s; 72 °C for 5 min; 98 °C for 1 min; and hold at 4 °C. PCR1 products were purified with AMPure XP beads, quantified using PicoGreen and diluted to 70 pg μl⁻¹.

PCR2 was used to add the remaining Illumina adapter sequences and sample indexes. PCR1 products were diluted to 7.2 × 10^5^ single-stranded DNA molecules per microlitre, so that the number of input molecules was lower than the anticipated sequencing depth, enabling generation of barcode families with multiple reads per UMI. Each PCR2 reaction contained 1 μl diluted PCR1 template, corresponding to 7.2 × 10^5^ single-stranded DNA molecules, and 2 μl each of universal Illumina forward primer and indexed reverse primer. Cycling conditions were: 98 °C for 30 s; 30 cycles of 98 °C for 10 s, 56 °C for 10 s and 72 °C for 10 s; 72 °C for 5 min; and hold at 4 °C. Final PCR products were purified with AMPure XP beads, analysed on a 1% agarose gel and quantified. Libraries of approximately 250 bp, depending on the sample, were pooled in equimolar amounts and sequenced by Genewiz on an Illumina HiSeq 2500 platform using 2×250 bp paired-end reads.

### Sequencing reads analyses

Paired-end 250-bp reads were processed into consensus sequences using the consensusmaker.py script^58^. Briefly, raw reads were grouped into families based on 16-nt barcodes generated by concatenating the 8-nucleotide barcode sequences present at each end of the reads. Reads sharing identical barcodes were collapsed into consensus sequences, each representing a single RNA molecule prior to PCR amplification, thereby removing PCR duplicates. Families were required to contain a minimum of 3 and a maximum of 1000 reads, and mutations were incorporated into the consensus only if present in at least 70% of reads within a family.

Consensus sequences were filtered to remove residual Illumina adapter sequences using CUTADAPT v4.3^59^. Cleaned reads were aligned to the reference genome using BWA-MEM v0.7.17^60^ with default parameters. The resulting SAM files were converted to BAM format and sorted using samtools v1.19^61^. To improve indel detection and mapping accuracy, BAM files were subsequently realigned using GATK v3.8^62^.

Variant positions were identified from the BAM files using samtools mpileup (maximum depth -d set to 500,000 reads per position). Although total sequencing depth ranged from 700,000 to 1,000,000 consensus sequences, this upper limit was applied to standardize coverage across samples. Mutation positions were extracted using the script mut-position.py^58^, resulting in a comprehensive table of detected variants available in our Zenodo repository along with the full analysis pipeline (link available upon request to the corresponding author).

### Insertion mapping and analysis

Insertion calls were obtained from the comprehensive table of detected variants, which provides, for each detected insertion, its attributed genomic position. For each position, insertion counts and frequencies were computed using insertion_counter.py, a custom Python script available at our Zenodo repository (link available upon request to the corresponding author), which outputs a workbook (insertions.xlsx) containing sequencing depth and counts of all observed insertion types.

Short insertions in homopolymeric regions can be ambiguously positioned during realignment with GATK v3.8^62^. For example, an AG insertion or a GA insertion in the parental sequence CAAAGAAAAAAAAGAGG can generate the same final sequence, CAAAGAGAAAAAAAAGAGG, depending on the coordinate to which the insertion is assigned (CAA**<u>AG</u>**AGAAAAAAAAGAGG or CAAA**<u>GA</u>**GAAAAAAAAGAGG). In these rare cases, ambiguous insertions were assigned to the coordinate closest to the local homopolymeric region that yielded an equivalent inserted sequence. In the illustrative example above, the insertion was therefore assigned to the GA insertion yielding CAAA**<u>GA</u>**GAAAAAAAAGAGG.

### Transient secondary RNA structure analyses

Predicted transient RNA structures were computed using the sliding window approach implemented in the slidingfold.py script from the repository of Mathis Funk^18^ available at GitHub (https://github.com/dr-funk/trapped-RdRp). The -t option was used to extract position-specific information, including predicted secondary structures and their associated stability values, for each sequence in tab-separated format.

The resulting output files are available at our Zenodo repository (link available upon request to the corresponding author).

### Stationary secondary RNA structure analyses

Stationary RNA secondary structures were predicted using RNAfold from the ViennaRNA package available at ViennaRNA Web Services^63^. An 80-nucleotide window encompassing the haemagglutinin cleavage site and its flanking sequences was used to capture the local nucleotide context. Minimum free energy (MFE) structures were computed using default parameters, and structural representations were derived directly from the RNAfold output. The list of 80-nt windows analysed for each virus is available in our Zenodo repository.

### Calculation of ΔG and ΔΔG values

To calculate thermodynamic stability differences between original and realigned product– template duplexes formed during viral RdRp slippage, we developed inserpredictor.py, a script that takes viral sequences in FASTA format as input. For each position and for both cRNA and vRNA synthesis, the script extracts the nine-nucleotide template sequence predicted to occupy the viral RdRp catalytic site and generates the corresponding nascent product sequence. The script is available at our Zenodo repository (link available upon request to the corresponding author).

The original product–template duplex was modelled as a fully complementary nine-base-pair RNA heteroduplex, with thermodynamic stability ΔGᵢ. Duplexes were represented using dot-bracket interaction strings, in which paired positions were indicated by parentheses and mismatched or unpaired positions by dots, with the two RNA strands separated by “&”. The stability of each duplex was calculated using the ViennaRNA Python module^63^, specifically the function RNA.eval_structure_simple.

To model viral RdRp slippage, the script calculated the stability of all putative product– template duplexes formed after downstream displacement of the nascent product RNA by one to nine nucleotides, yielding ΔG₁ to ΔG₉. These realigned duplexes may contain mismatches and therefore differ in stability from the original duplex. For each displacement n, the stability difference was calculated as ΔΔG = ΔGₙ - ΔGᵢ, where ΔGₙ corresponds to the realigned product–template duplex formed after an n-nucleotide displacement. All ΔG and ΔΔG values for candidate slippage events on cRNA and vRNA templates were stored in the crnaDDG.xlsx and vrnaDDG.xlsx files, respectively. These workbooks are the output of inserpredictor.py and are available in our Zenodo repository (link available upon request to the corresponding author).

### Classification of 3′ terminal match or mismatch

Upon viral RdRp slippage, the 3′ end of the reannealed product–template duplex may be correctly base-paired with the template or may form a 3′ terminal mismatch. Previous work has shown that the identity of the 3′ terminal base pair or mismatch differentially influences elongation from reannealed duplexes, with some mismatches being tolerated more efficiently than others^34–36^. We therefore encoded the 3′ terminal interaction as a categorical elongation parameter rather than as a simple match/mismatch variable. Based on these experimentally defined mismatch-tolerance categories, and our observation of enhanced insertion for A:U or U:A 3′ product matches in the presence of an upstream polyadenine or polyuracil tract, we assigned the following categories: MA for A:U or U:A terminal matches within polyadenine or polyuracil tracts; MB for G:C, C:G, U:A, A:U and G:U matches or wobble pairs; MC for tolerated terminal mismatches, C:A, C:U and U:U; and MD for poorly tolerated terminal mismatches, G:A, G:G and C:C.

### Thermodynamic pathway accessibility

For each candidate slippage event, inserpredictor.py evaluated the ΔG and ΔΔG values of all intermediate product–template duplexes encountered between the original duplex and the final realigned duplex. Because slippage to distal positions requires sequential progression through these intermediate states, a predicted insertion was considered accessible only if the full slippage trajectory remained thermodynamically permissive. Slippage trajectories were classified according to displacement length as short (<4 nucleotides) or long (≥4 nucleotides). For each class, ΔG and ΔΔG accessibility thresholds were calibrated during model development using only the H5-Shim training series, by comparing experimentally observed insertion profiles with predictions generated before applying the thermodynamic pathway-accessibility criterion. Thresholds were selected to exclude predicted false-positive insertions, including short and long insertions not detected experimentally, while retaining experimentally observed insertions. The exact ΔG and ΔΔG thresholds used to classify short and long slippage trajectories as accessible are provided in table S3. These fixed thresholds were then applied without modification to all validation, natural-emergence and database analyses. Insertions failing this accessibility criterion were excluded from *HPAIVpredict* output.

### Training of the *HPAIVpredict* model and predictions

Predictor variables were extracted from the crnaDDG.xlsx and vrnaDDG.xlsx output files generated by the ΔG calculation module. For each candidate slippage event, the model used two predictors: (i) the thermodynamic stability difference between the original and realigned product–template duplexes, ΔΔG = ΔGₙ − ΔGᵢ, and (ii) the 3′ terminal interaction category of the reannealed duplex, classified as MA, MB, MC or MD.

The response variable used for model training was derived independently from experimentally observed insertion frequencies measured by single-strand consensus sequencing. Because frequency estimates for lower-abundance insertions are expected to be affected by higher relative sampling uncertainty, model training was restricted to experimentally detected insertions with frequencies ≥ 10⁻⁴ to reduce the influence of this uncertainty on coefficient estimation. For these experimentally detected insertions, the corresponding ΔΔG value and 3′ terminal interaction category were retrieved from the ΔG output files, and the experimentally observed insertion frequency was used as the response variable after log₁₀ transformation.

Insertion frequencies were modelled using a multiple linear regression implemented in R with the lm function. The model was trained on experimentally observed insertions from the H5-Shim variants (Shim, ShimA1, ShimA2, ShimA3, ShimA4, Shim24a2b) and the PR8 H1 sequence. The regression model was defined as:

log_10_(f)=β0+α*(ΔΔG)+ηM+ε

where f is the experimentally observed insertion frequency, β_0_ is the intercept, α is the fitted coefficient associated with ΔΔG, ηM is the fitted category-specific term associated with the 3′ terminal interaction category M, and ε is the residual error term.

Following model training, the fitted parameters were: β_0_ =-1.10587, α=-0.31829, ηMB=-1.46627, ηMC=-1.60555 and ηMD=-2.87328. MA was used as the reference category during model fitting and therefore ηMA= 0.

Predicted log_10_ insertion frequencies for all candidate slippage events were then calculated using the R predict.lm function from their ΔΔG values and 3′ terminal match/mismatch categories. Predicted insertions with frequencies below the sequencing-based detection threshold of 2×10^-6^ per position were excluded from downstream *HPAIVpredict* outputs. The remaining predicted insertions were subsequently filtered using the thermodynamic pathway-accessibility criterion described above. This filtering step excluded predicted insertions whose final realigned duplex displayed favourable ΔΔG values but whose slippage trajectory required traversal of one or more unstable intermediate product-template duplexes. Such trajectories were considered thermodynamically inaccessible because unstable intermediates would be expected to act as barriers to progression along the slippage pathway. Thus, *HPAIVpredict* estimates insertion frequencies from ΔΔG and the 3′ terminal match/mismatch category using a regression model, and retains predicted insertions only when the corresponding slippage trajectory satisfies an independent thermodynamic pathway-accessibility criterion.

All steps were implemented in *HPAIVpredict*, a custom pipeline available at our Zenodo repository (link available upon request to the corresponding author).

### H5 and H7 database construction and predictions of insertions and furin cleavability

HA sequences were retrieved from the GISAID EpiFlu database on 4 November 2025 using the Search & Browse interface. All H5 (2,814) and H7 (5,580) sequences annotated as low pathogenic avian influenza viruses (LPAIV) were selected across all available geographic regions. Default settings were used for all other search parameters. Duplicate sequences were removed using seqkit rmdup -s, and duplicate sequence identifiers were removed using seqkit rmdup without the -s option.

To exclude residual highly pathogenic sequences, multiple sequence alignments were generated using Clustal Omega v1.2.4 with default parameters, and sequences containing insertions at the haemagglutinin cleavage site relative to subtype-specific LPAIV consensus motifs were removed.

The curated dataset containing 2191 unique H5 sequences and 3287 unique H7 sequences was processed using a custom Python script. For each sequence, a 44-nucleotide window centred on the haemagglutinin cleavage site was extracted, including flanking nucleotides on both the 3′ and 5′ sides of the cleavage site-encoding region, to capture sequence contexts potentially engaged during vRNA- and cRNA-synthesis slippage around the central cleavage-site sequence. Sequences sharing an identical 44-nt window were grouped, and each group was collapsed into a single representative sequence, with headers retaining the identifiers of all grouped sequences. This grouping yielded 344 H5 and 286 H7 representative 44-nucleotide sequence contexts.

For each unique 44-nt sequence, in silico mutagenesis was performed by systematically introducing adenine substitutions along the central cleavage site window. This generated independent sequence sets, corresponding to the original sequence (“WT” sheet in the respective H5 and H7 workbooks), individual substitutions (Mut1, Mut2 and Mut3), and the most relevant combinations of substitutions based on experimental results identifying a prominent role of specific combinations of substitutions in H5 and H7, respectively. Each dataset was transformed into FASTA files for subsequent analysis.

Each FASTA set was individually analysed using HPAIVpredict v1.0.0 to predict nucleotide insertion events in the HA cleavage site-encoding sequence. The resulting predictions were merged and converted into a tabular format using a custom R script. Each table contained one representative sequence per line and, for each position in the HA cleavage site-encoding sequence, the “cumulative insertion frequency” of all predicted insertion events, expressed as log_10_-transformed insertion frequency (data S1 and data S2).

For insertions with lengths that were multiples of three nucleotides, corresponding to in-frame insertions, the resulting nucleotide sequences were reconstructed and translated into amino acid sequences. The amino acid sequences resulting from in-frame insertions were recorded in the output table at the corresponding insertion positions. All in-frame insertions were analysed for predicted furin-dependent cleavability using ProP v1.0^45^ to generate predicted furin-cleavage scores using the ProP online server with the “generate plot” and “general PC prediction” options disabled and the “verbose” option enabled. HA cleavage-site amino acid sequences yielding ProP furin-cleavability scores equal to or greater than those of naturally occurring H5 (score ≥ 0.46) or H7 (score ≥ 0.19) HPAIV cleavage sites were classified as predicted furin-cleavable MBCS and recorded in the “predicted furin-cleavable HA cleavage site (ProP cleavability score)” line with the corresponding amino acid sequences highlighted in blue and their ProP cleavability scores indicated in brackets (data S1 and data S2).

### Precursor H5 and H7 sequence inference

When unknown, LPAIV precursor sequences of H5 and H7 HPAIV were generated by replacing the multibasic cleavage site (MBCS) of each HPAIV with subtype-specific canonical monobasic cleavage site sequences. This approach preserves the genetic background of the corresponding HPAIV while approximating a low-pathogenic ancestor that differs primarily at the cleavage site. This strategy is supported by documented emergence events in which HPAIV HAs differ minimally from those of their LPAIV precursors outside the cleavage site.

## Data and materials availability

Raw next-generation sequencing data generated in this study have been deposited in the NCBI Sequence Read Archive under BioProject accession PRJNA1481509. Individual SRA run accessions, sample identifiers and associated metadata are provided in data S3. Processed data supporting the figures and analyses are provided in data S4. The bioinformatics pipeline, analysis scripts and code used for read processing, quality control, mapping, variant calling, downstream analysis and figure generation will be made available upon request to the corresponding author and made publicly available upon publication.

## Supplementary Materials

**Fig. S1.**
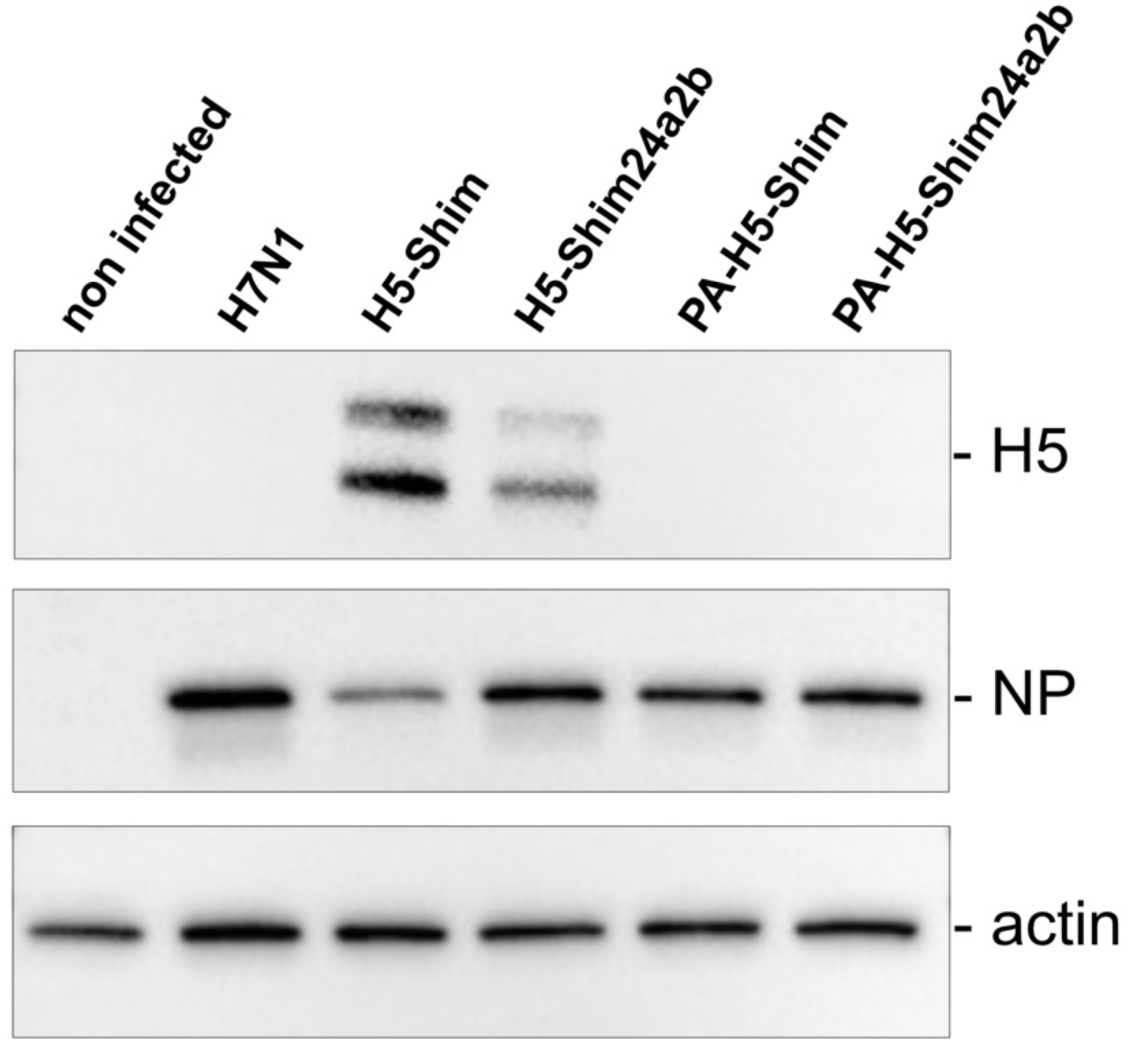
H5 HA protein is not detected in cells infected with PA-H5-Shim or PA-H5-Shim24a2b viruses. Western blot analysis of protein expression in MDCK cells infected with recombinant influenza viruses. Cells were either left uninfected or infected with wild-type A/turkey/Italy/977/1999 (H7N1), recombinant viruses expressing functional H5 HA from A/whistling swan/Shimane/499/83 (H5-Shim) or its Shim24a2b variant (H5-Shim24a2b), or PA-H5-Shim and PA-H5-Shim24a2b viruses carrying the corresponding H5 sequences as non-translatable transgenes. H5 HA was detected in cells infected with viruses expressing functional H5-Shim or H5-Shim24a2b, but not in cells infected with PA-H5-Shim or PA-H5-Shim24a2b. Viral NP detection confirms influenza virus infection, and actin serves as a loading control.

**Fig. S2.**
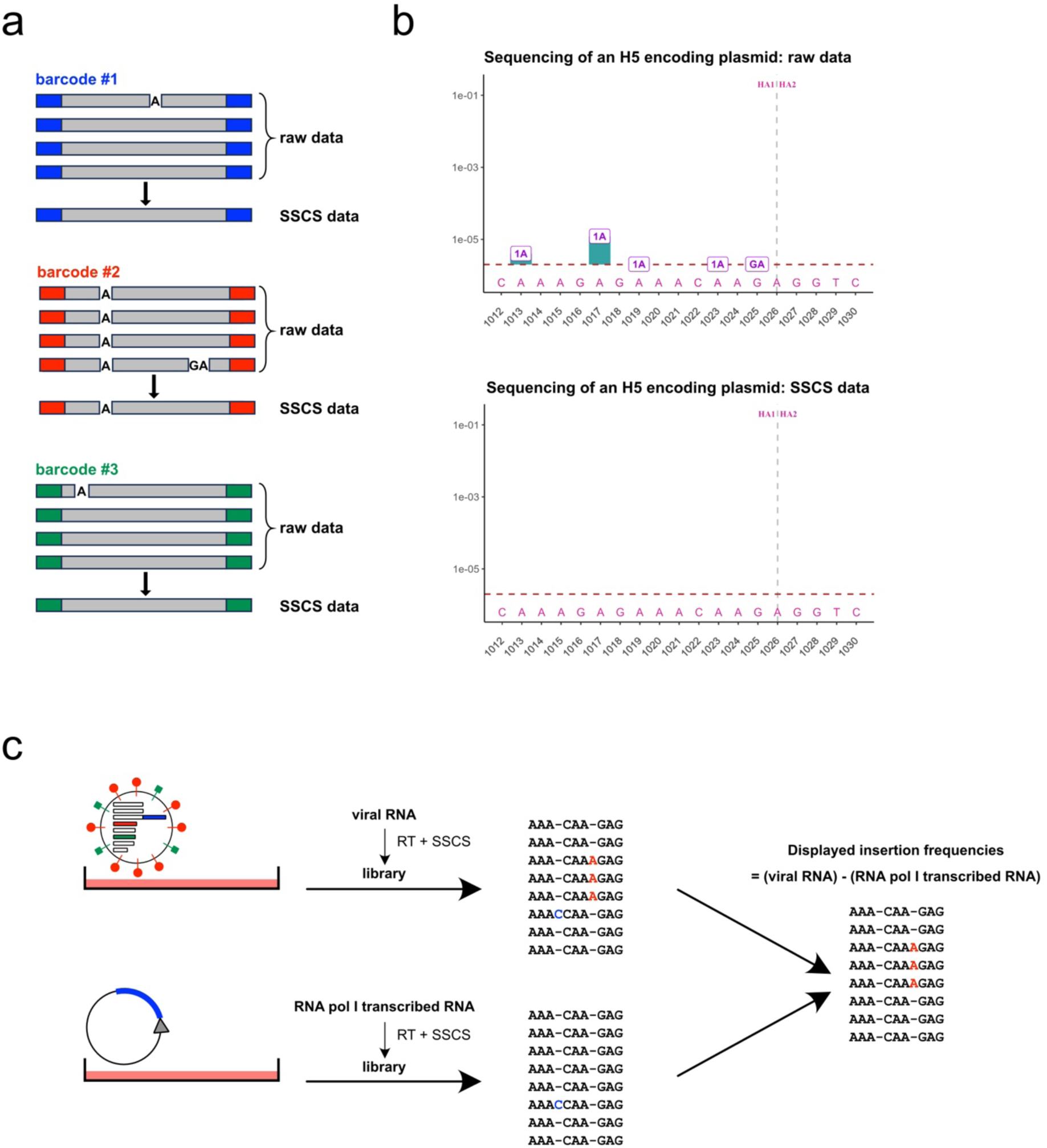
Description of the workflow used to accurately detect viral RdRp-mediated insertions. a) Principles of the Illumina-based single strand-consensus sequencing method. A first round of PCR appends random barcodes and part of the Illumina adapter to each amplicon. Sequencing reads are grouped by barcode, distinguishing sequencing errors that occur in only one read (raw data) from true insertions that occur in all reads and that are thus contained in single strand-consensus sequencing (SSCS) data. b) Analysis of sequencing of an H5-encoding plasmid. For clarity, only insertions in a 19-nucleotide window corresponding to nucleotides 1012 to 1030 of the H5 cleavage site are represented in the graphs. The histogram bars represent the average frequency of insertions at each position calculated from two independent experiments. The letters indicate the nucleotide composition of each type of insertion at the indicated position in the H5 cleavage site sequence, while their position along the y-axis corresponds to the frequency of each type of insertion. This analysis reveals that raw data contains insertions that correspond to PCR or sequencing errors, which are absent from SSCS data from a 2 × 10⁻⁶ threshold, indicated by the dotted line. c) Strategy used to correct for reverse-transcriptase (RT)-mediated errors and for errors potentially introduced by the human RNA polymerase I (RNA pol I) during the virus rescue process. For each HA sample tested in the PA-HA system, we transfect the cells in parallel culture dishes with plasmid expressing the corresponding HA transcribed from a human RNA polymerase I promoter. RNA collected from these parallel samples was subjected to RT and SSCS. We subtract nucleotide insertions detected in parallel control experiments from insertions detected in the PA-HA samples to accurately report on the identity and frequencies of nucleotide insertions caused by the viral RdRp.

**Fig. S3.**
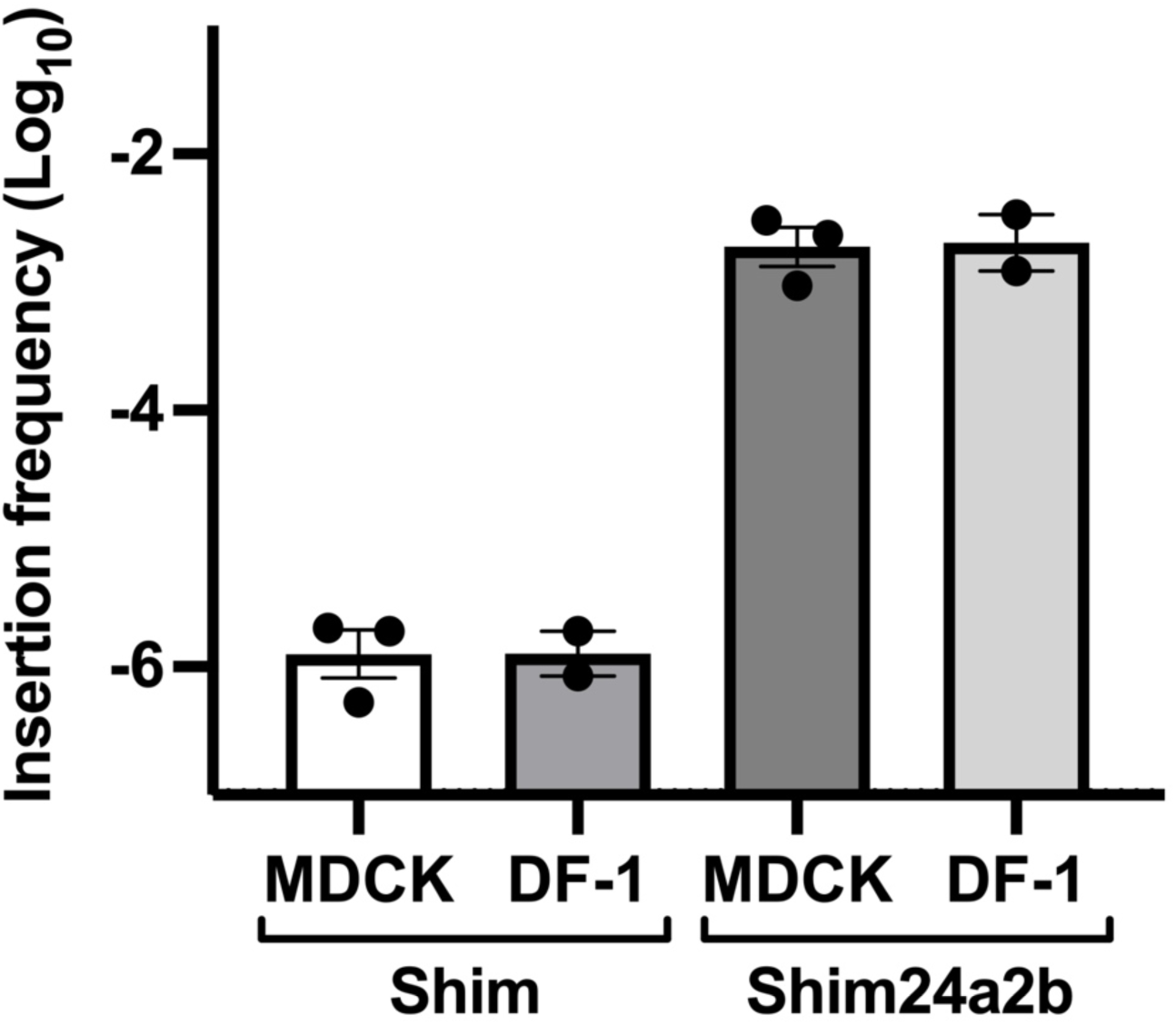
Insertion frequencies in canine MDCK and in chicken DF-1 cells. Quantification of insertion events across all lengths in the HA cleavage sites of PA-H5-Shim and PA-H5-Shim24a2b grown either in canine MDCK or in chicken DF-1 cells. For each independent experiment, insertion frequencies were averaged across the 19-nucleotide window. Bars show mean ± s.e.m.; points represent independent experiments.

**Fig. S4.**
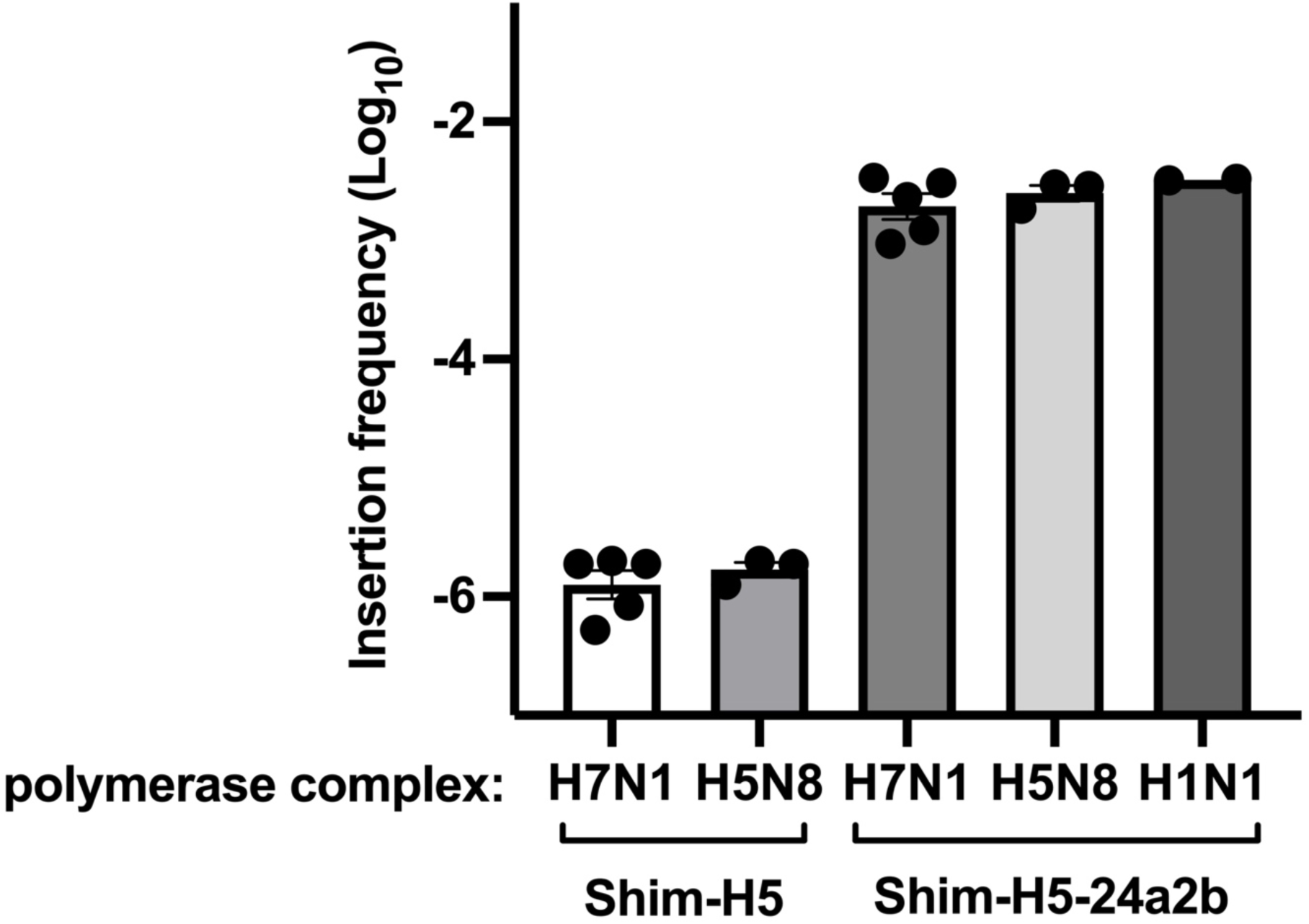
Insertion frequencies with different influenza A viral RdRp complexes. Quantification of insertion events across all lengths in the Shim and Shim24a2b HA cleavage-site sequences. Recombinant viruses carrying PA-H5-Shim or PA-H5-Shim24a2b transgenes and expressing either the H7N1 RdRp complex of A/turkey/Italy/977/1999 (H7N1) or the H5N8 HPAIV RdRp complex of A/mallard duck/France/171201g/2017 (H5N8) were grown in MDCK cells. In parallel, minigenome assays were performed in human 293T cells using the Shim24a2b HA sequence as the viral RNA substrate and the H1N1 RdRp complex of A/WSN/1933 (H1N1). For each independent experiment, insertion frequencies were averaged across the 19-nucleotide window spanning the HA cleavage-site–encoding sequence. Bars show mean ± s.e.m.; points represent independent experiments.

**Fig. S5.**
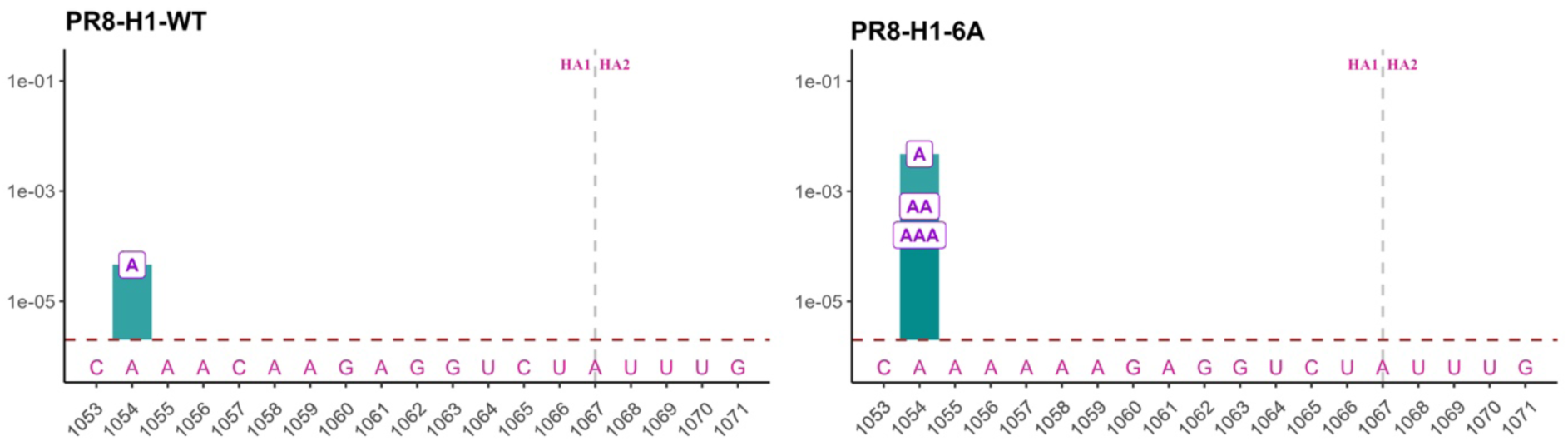
A six-adenine stretch converts PR8 H1 to an insertion-prone sequence. Nucleotide insertion profiles in the HA cleavage site encoding sequence of A/Puerto Rico/8/1934 (H1N1) (PR8). Wild-type PR8 HA (PR8-H1-WT) was compared to PR8-H1-6A carrying a six-adenine stretch created by a C1016A substitution. Insertions in a 19-nucleotide window corresponding to nucleotides 1012 to 1030 of the H1 cleavage site are represented in the graphs. The histogram bars represent the insertion frequencies at each position. The dotted horizontal line corresponds to the 2 × 10⁻⁶ sequencing detection threshold. The vertical dotted line indicates the site of HA cleavage.

**Fig. S6.**
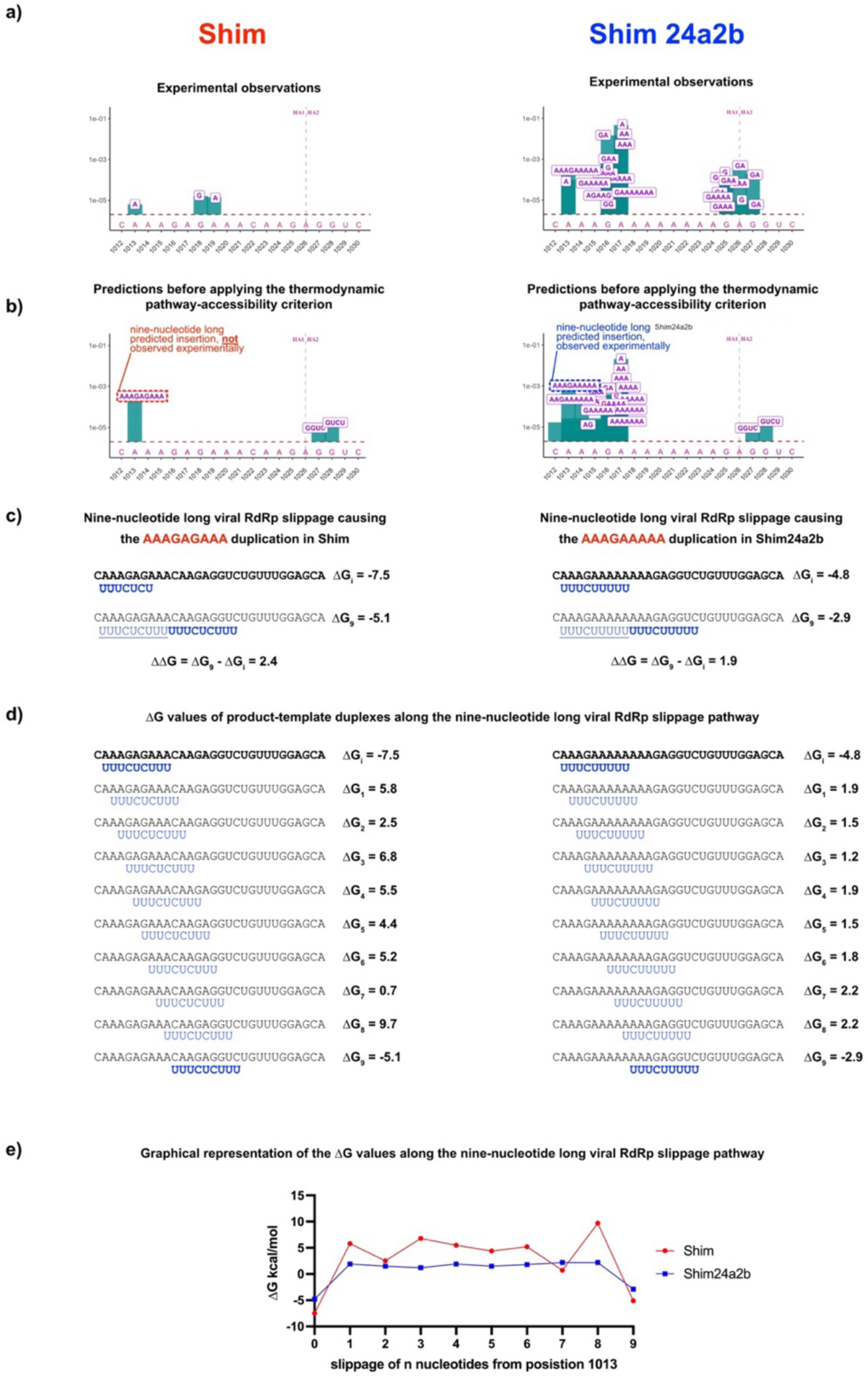
Illustration of the thermodynamic pathway accessibility for a nine-nucleotide long viral RdRp slippage highlighting unstable intermediate duplexes present in H5-Shim but absent in H5-Shim24a2b. Left panels show PA-H5-Shim; right panels show PA-H5-Shim24a2b. **a)** Experimentally observed insertion profiles within a 19-nucleotide window spanning positions 1012–1030 of the H5 HA cleavage-site–encoding sequence. **b)** Insertions predicted by the two-parameter model before applying the thermodynamic pathway-accessibility criterion. A predicted nine-nucleotide insertion is highlighted; this insertion is not observed experimentally in PA-H5-Shim but is observed in PA-H5-Shim24a2b. **c)** Thermodynamic comparison of the predicted nine-nucleotide RdRp slippage leading to duplication of the indicated template sequence. The final ΔΔG values are similar for PA-H5-Shim and PA-H5-Shim24a2b, explaining why this insertion is predicted in both sequence contexts before incorporation of the thermodynamic pathway-accessibility criterion. **d)** ΔG values of successive intermediate product–template duplexes along the nine-nucleotide slippage pathway. **e)** Graphical representation of the ΔG values shown in d. In PA-H5-Shim, the slippage pathway is interrupted by intermediate duplexes with high, unfavourable ΔG values, suggesting a thermodynamic barrier to progression towards the final backtracked duplex. By contrast, intermediate ΔG values remain more favourable in PA-H5-Shim24a2b, consistent with the experimentally observed nine-nucleotide insertion. Thus, the thermodynamic pathway-accessibility criterion improves model specificity by excluding predicted insertions that require traversal of unfavourable intermediate duplexes.

**Fig. S7.**
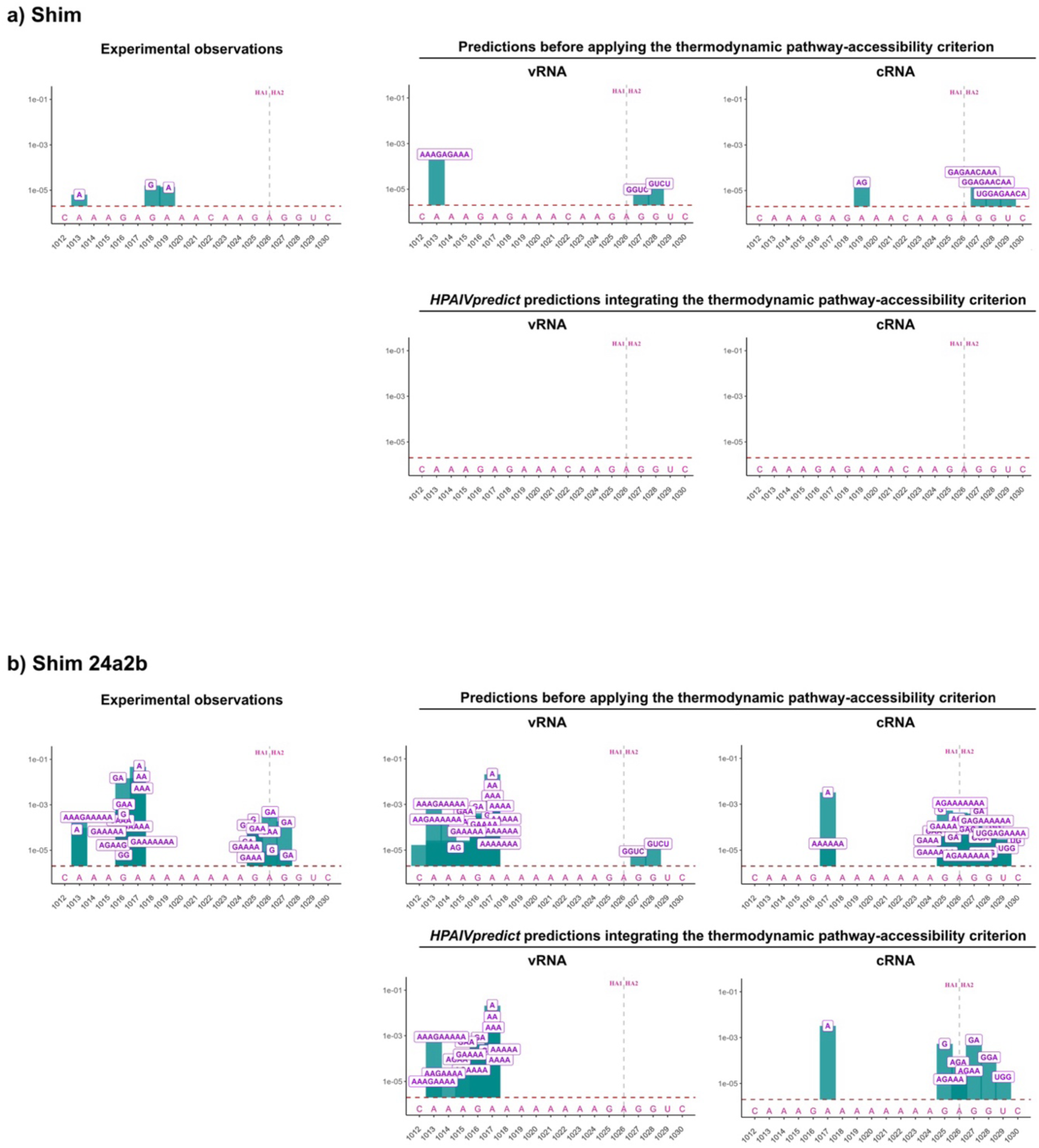
Observed insertion profiles in the H5 cleavage site of Shim and Shim24a2b and predicted insertions before and after applying the thermodynamic pathway-accessibility criterion. Comparison of experimentally observed insertions (left) and insertions predicted before thermodynamic pathway-accessibility criterion was applied (top) or predictions integrating the thermodynamic pathway-accessibility criterion (bottom) in a 19-nucleotide window corresponding to nucleotides 1012 to 1030 of the H5 cleavage site of Shim (a) and Shim24a2b (b). The histogram bars for observed insertions represent the average frequency of insertions at each position calculated from five independent experiments. The letters indicate the nucleotide composition of each type of insertion at the indicated position in the H5 cleavage site sequence, while their position along the y-axis corresponds to the frequency of each type of insertion. The dotted horizontal line corresponds to the 2 × 10⁻⁶ sequencing detection threshold. The vertical dotted line indicates the site of HA cleavage.

**Fig. S8.**
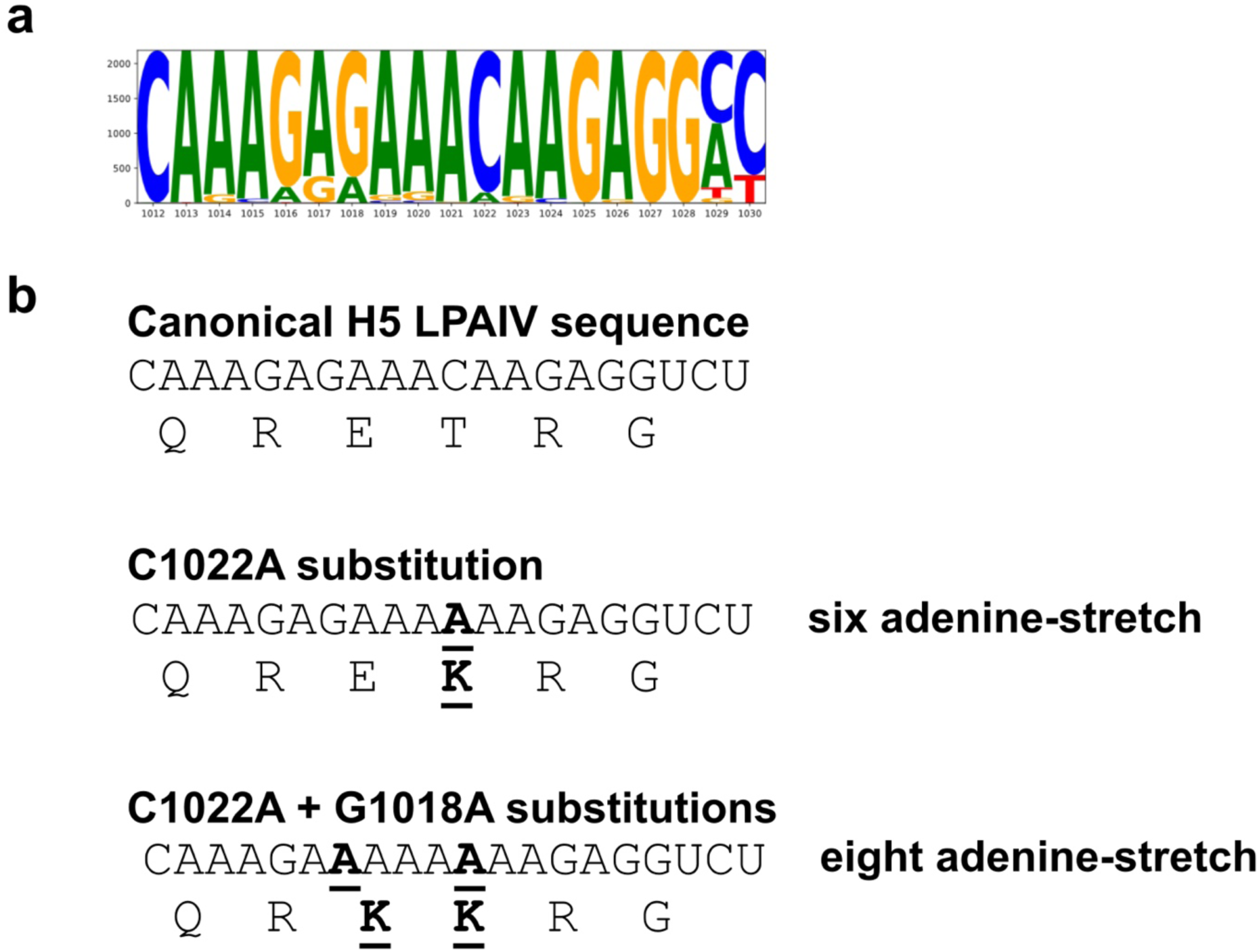
H5 LPAIV HA cleavage-site sequences and generation of adenine tracts by substitution. **a)** Sequence logo of the HA cleavage-site–encoding region (positions 1012–1030) derived from all H5 low-pathogenic avian influenza viruses (LPAIVs) available in the GISAID database. Letter height reflects nucleotide frequency at each position, highlighting the canonical sequence context and the general absence of extended adenine tracts in circulating H5 LPAIVs. **b)** Representative H5 HA cleavage-site sequences illustrating the stepwise generation of adenine stretches through point mutations. Top, canonical LPAIV sequence encoding a monobasic cleavage site (RETR). Middle, a single C1022A substitution generates a six-adenine tract and a dibasic cleavage site (REKR). Bottom, combination of C1022A and G1018A substitutions produces an eight-adenine stretch and RKKR cleavage-site motif. Substituted nucleotides are indicated, and corresponding amino acid translations are shown below each sequence.

**Fig. S9.**
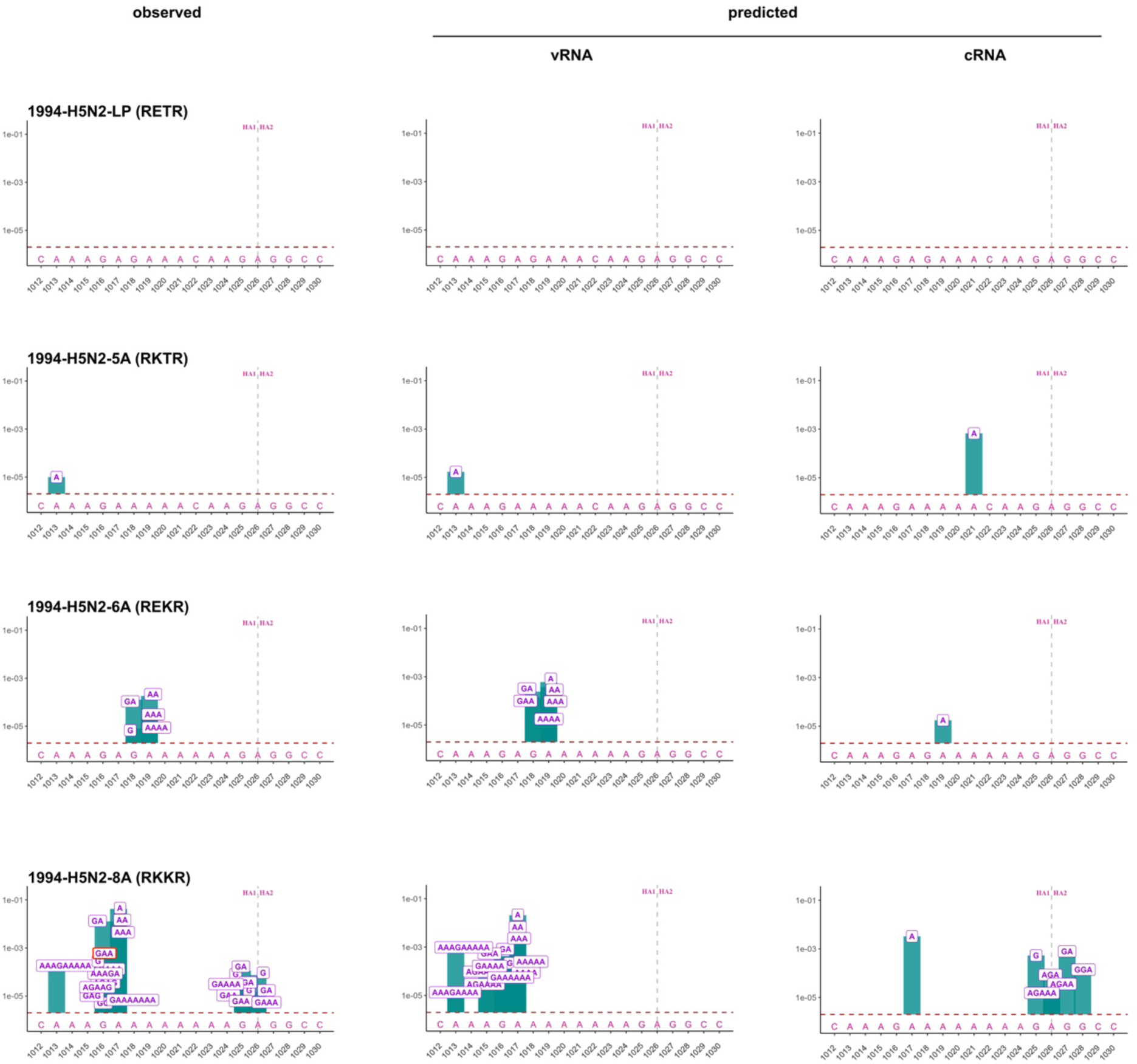
Insertion profile in adenine-rich intermediates of the 1994-H5N2 emergence variants. Comparison of experimentally observed insertions (left) and insertions predicted to occur during vRNA and cRNA synthesis (centre and right, respectively) for the 1994-H5N2 emergence event. The canonical LPAIV precursor (RETR cleavage site), a five-adenine variant (5A-RKTR), a six-adenine variant (6A-REKR) and an eight-adenine variant (8A-RKKR) were analysed. The histogram bars represent the average frequency of insertions at each position calculated from two to five independent experiments. The letters indicate the nucleotide composition of each type of insertion at the indicated position in the H5 cleavage site sequence, while their position along the y-axis corresponds to the frequency of each type of insertion. The dotted horizontal line corresponds to the 2 × 10⁻⁶ sequencing detection threshold. The vertical dotted line indicates the site of HA cleavage. The GAA triplet insertion generating an elongated MBCS encoding the RRKKR motif found in several H5 HPAIVs is circled in red.

**Fig. S10.**
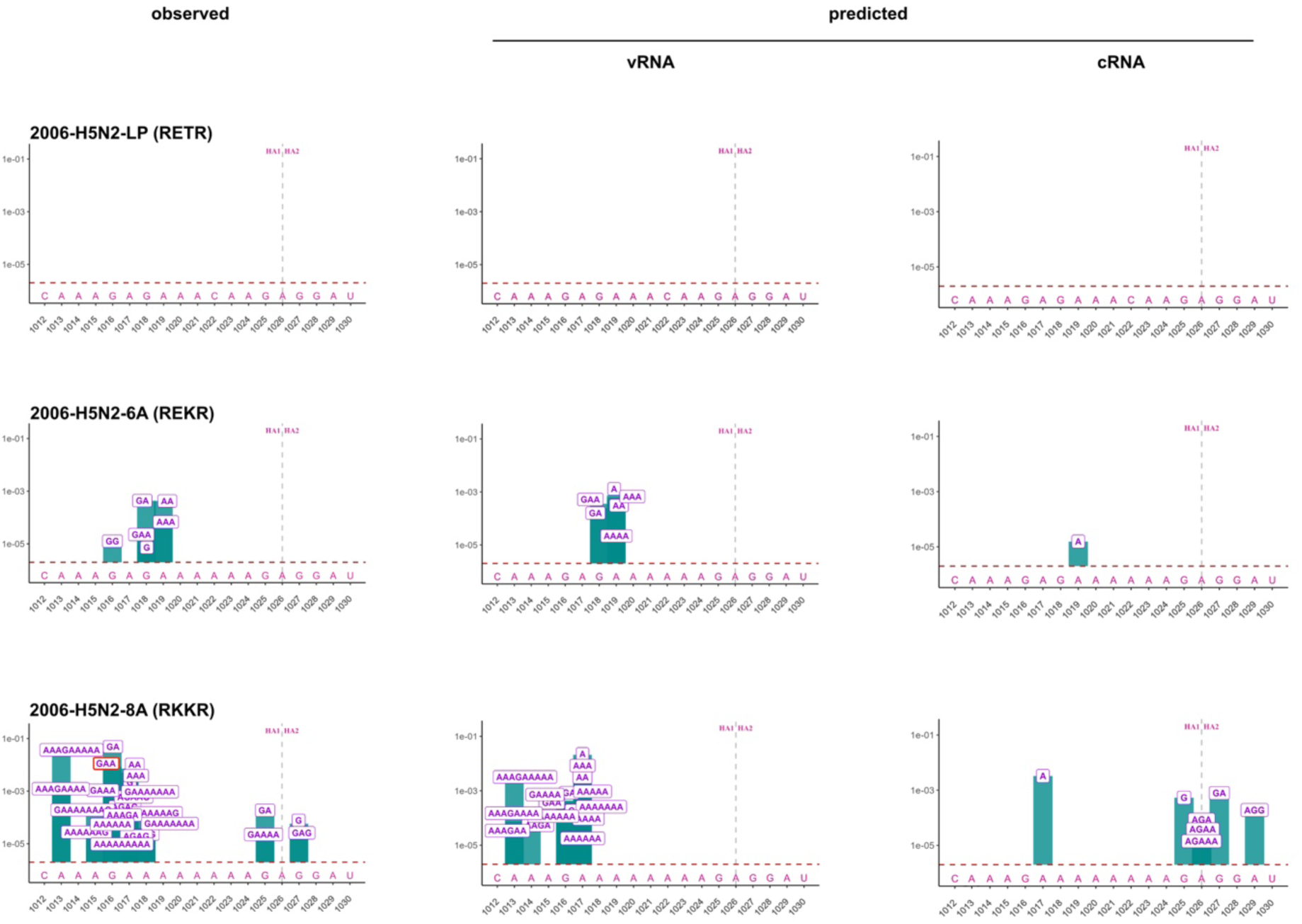
Insertion profile in adenine-rich intermediates of the 2006-H5N2 emergence variants. Comparison of experimentally observed insertions (left) and insertions predicted to occur during vRNA and cRNA synthesis (centre and right, respectively) for the 2006-H5N2 emergence event. The canonical LPAIV precursor (RETR cleavage site), a six-adenine variant (6A-REKR) and an eight-adenine variant (8A-RKKR) were analysed. The histogram bars represent the average frequency of insertions at each position calculated from two to five independent experiments. The letters indicate the nucleotide composition of each type of insertion at the indicated position in the H5 cleavage site sequence, while their position along the y-axis corresponds to the frequency of each type of insertion. The dotted horizontal line corresponds to the 2 × 10⁻⁶ sequencing detection threshold. The vertical dotted line indicates the site of HA cleavage. The GAA triplet insertion generating an elongated MBCS encoding the RRKKR motif found in several H5 HPAIVs is circled in red.

**Fig. S11.**
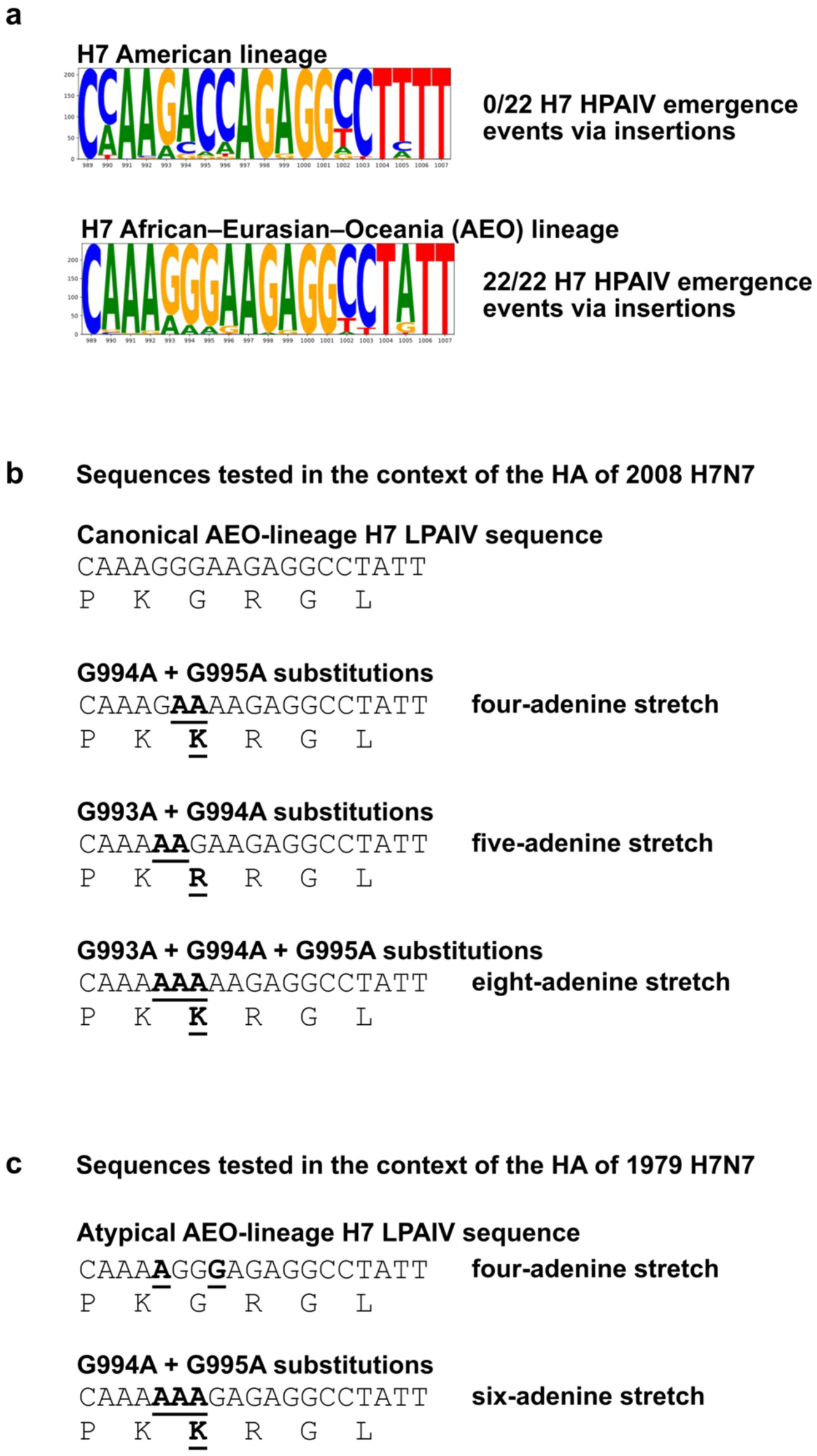
H7 LPAIV HA cleavage-site sequences and generation of adenine tracts by substitution. **a)** Sequence logo of the HA cleavage-site–encoding region (positions 980–1007) derived from all H7 low-pathogenic avian influenza viruses (LPAIVs) available in the GISAID database. According to their geographical origin, H7 LPAIVs were identified as belonging to the American lineage or to the African–Eurasian–Oceania (AEO) lineage. All H7 viruses that evolved to HPAIV via insertions belonged to the AEO lineage. Letter height reflects nucleotide frequency at each position, highlighting the canonical sequence context and the general absence of extended adenine tracts in circulating H7 LPAIVs. **b)** Representative H7 HA cleavage-site sequences experimentally tested in the context of the HA of 2008 H7N7. First row, canonical LPAIV sequence of the AEO lineage with a G_993_G_994_G_995_ triplet and typical H7 monobasic cleavage site (PKGR). Second row, combination of G994A and G995A substitutions in the canonical LPAIV sequence of the AEO lineage generates a four-adenine tract and PKKR cleavage site. Third row, combination of G993A and G994A substitutions in the canonical LPAIV sequence of the AEO lineage generates a five-adenine tract and PKRR cleavage site. Fourth row, combination of G993A, G994A and G995A substitutions in the canonical LPAIV sequence of the AEO lineage generates an eight-adenine tract and PKKR cleavage site. **c)** H7 HA cleavage-site sequences experimentally tested in the context of the HA of 1979 H7N7. Top row, atypical AEO-lineage H7 LPAIV sequence of the potential precursor of the 1979 H7N7 HPAIV. This LPAIV has a G993A substitution and an A996G in comparison with the canonical LPAIV sequence of the AEO lineage, generating a four-adenine tract, while maintaining a typical H7 monobasic cleavage site (PKGR). Bottom row: In this context, combined G994A and G995A substitutions create a six-adenine tract and PKKR cleavage site.

**Fig. S12.**
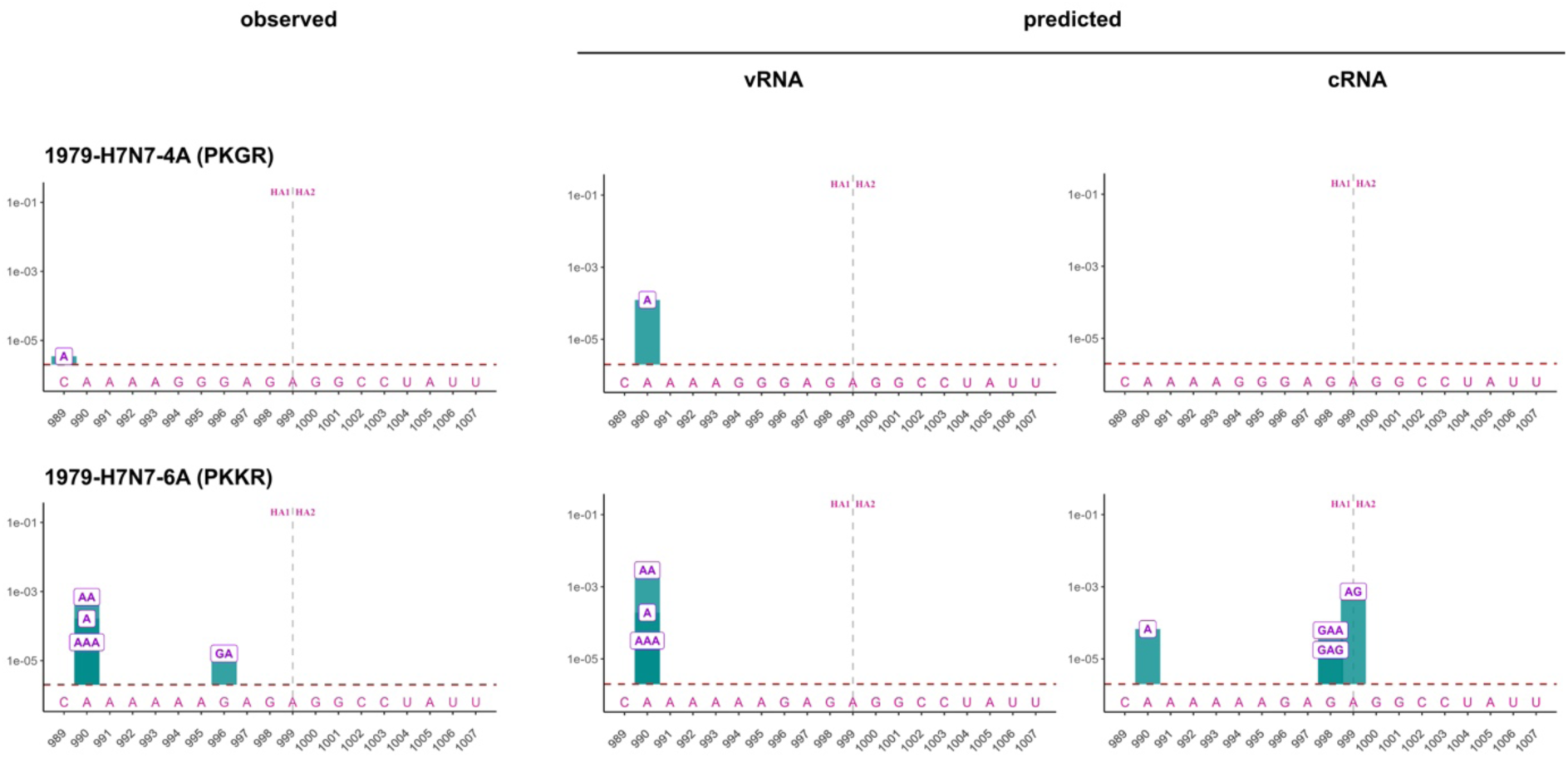
Insertion profile in adenine-rich intermediates of the 1979-H7N7 emergence variants. Comparison of experimentally observed insertions (left) and insertions predicted to occur during vRNA and cRNA synthesis (centre and right, respectively) for the 1979-H7N7 emergence event. The precursor with an atypical 4A-PKGR cleavage site and a six-adenine variant (6A-PKKR) were analysed. The histogram bars represent the average frequency of insertions at each position calculated from three independent experiments. The letters indicate the nucleotide composition of each type of insertion at the indicated position in the H7 cleavage site sequence, while their position along the y-axis corresponds to the frequency of each type of insertion. The dotted horizontal line corresponds to the 2 × 10⁻⁶ sequencing detection threshold. The vertical dotted line indicates the site of HA cleavage.

**Fig. S13.**
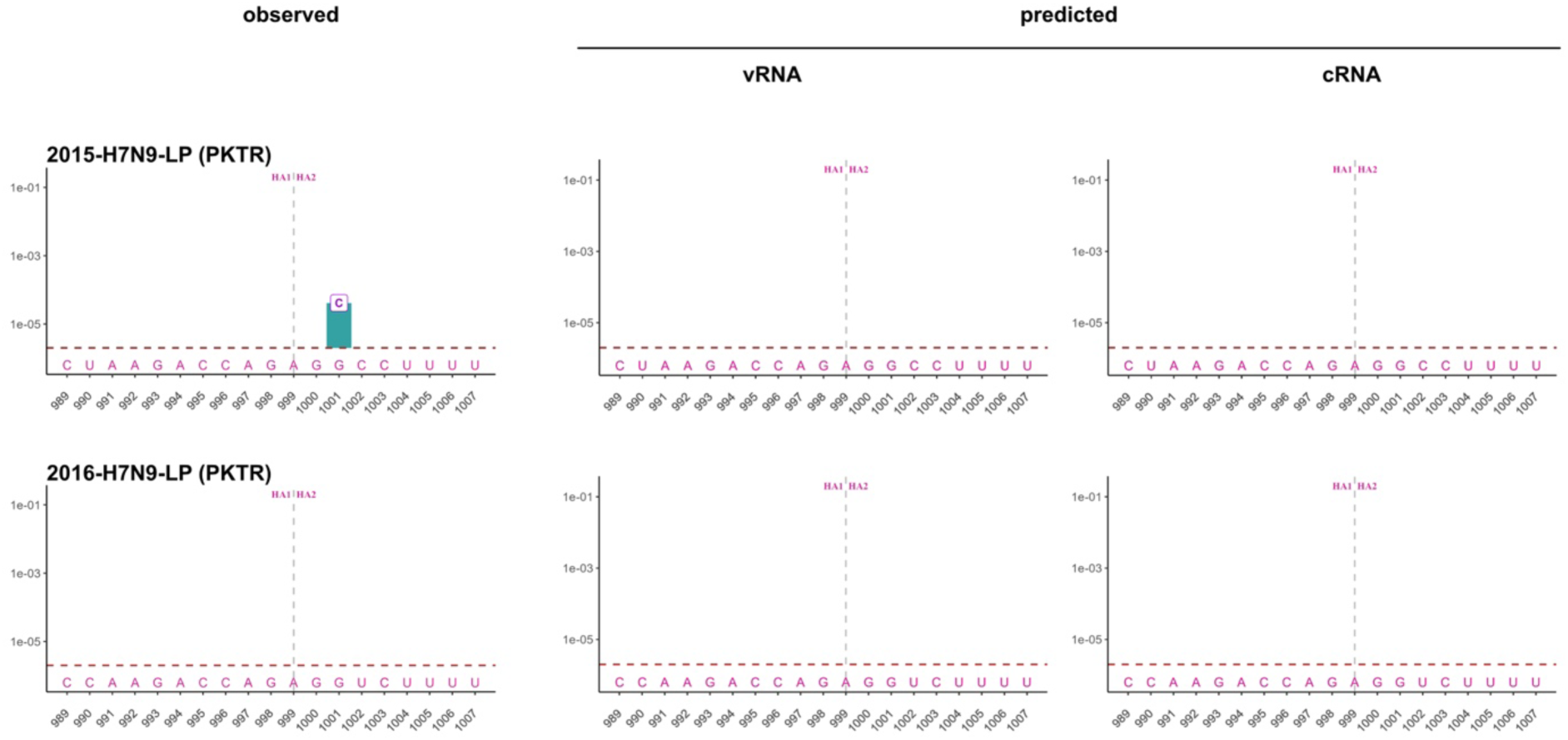
Insertion profile in two American-lineage H7 LPAIVs. Comparison of experimentally observed insertions (left) and insertions predicted to occur during vRNA and cRNA synthesis (centre and right, respectively) for two American-lineage H7 LPAIVs with a PKTR cleavage site. The 2015-H7N9-LP and the 2016-H7N9-LP were analysed. The histogram bars represent the average frequency of insertions at each position calculated from three independent experiments. The letters indicate the nucleotide composition of each type of insertion at the indicated position in the H7 cleavage site sequence, while their position along the y-axis corresponds to the frequency of each type of insertion. The dotted horizontal line corresponds to the 2 × 10⁻⁶ sequencing detection threshold. The vertical dotted line indicates the site of HA cleavage.

**Fig. S14.**
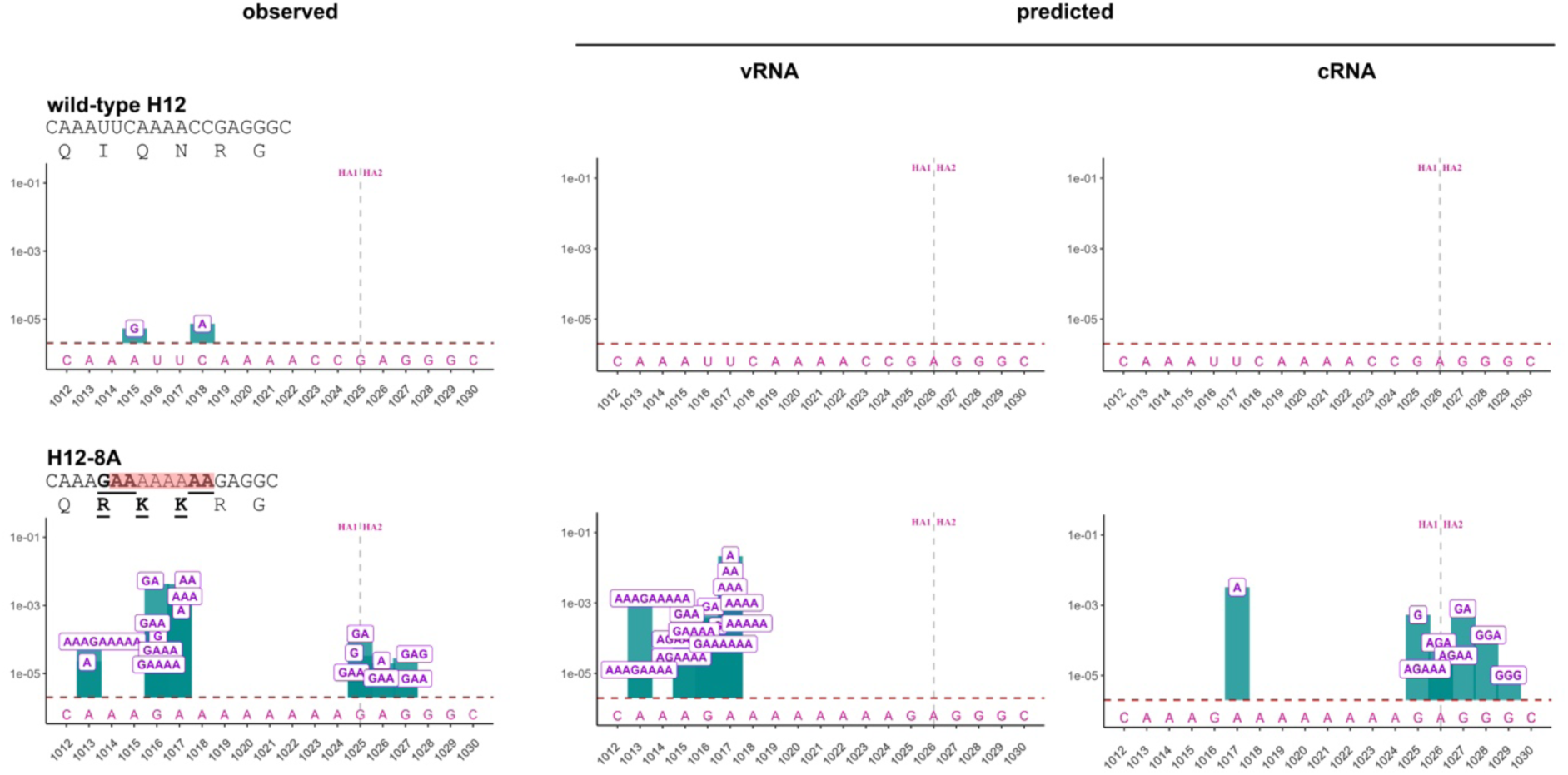
Five substitutions render a typical H12 insertion-prone. Comparison of experimentally observed insertions (left) and insertions predicted to occur during vRNA and cRNA synthesis (centre and right, respectively) in the HA cleavage site encoding sequence of A/duck/MW04/Vic/2012 (H12N5). Wild-type H12 was compared to H12-8A obtained following five substitutions indicated in bold and underlined. Insertions in a 19-nucleotide window corresponding to nucleotides 1012 to 1030 of the H12 cleavage site are represented in the graphs. The histogram bars represent the insertion frequencies at each position calculated from one experiment. The dotted horizontal line corresponds to the 2 × 10⁻⁶ sequencing detection threshold. The vertical dotted line indicates the site of HA cleavage.

**Fig. S15.**
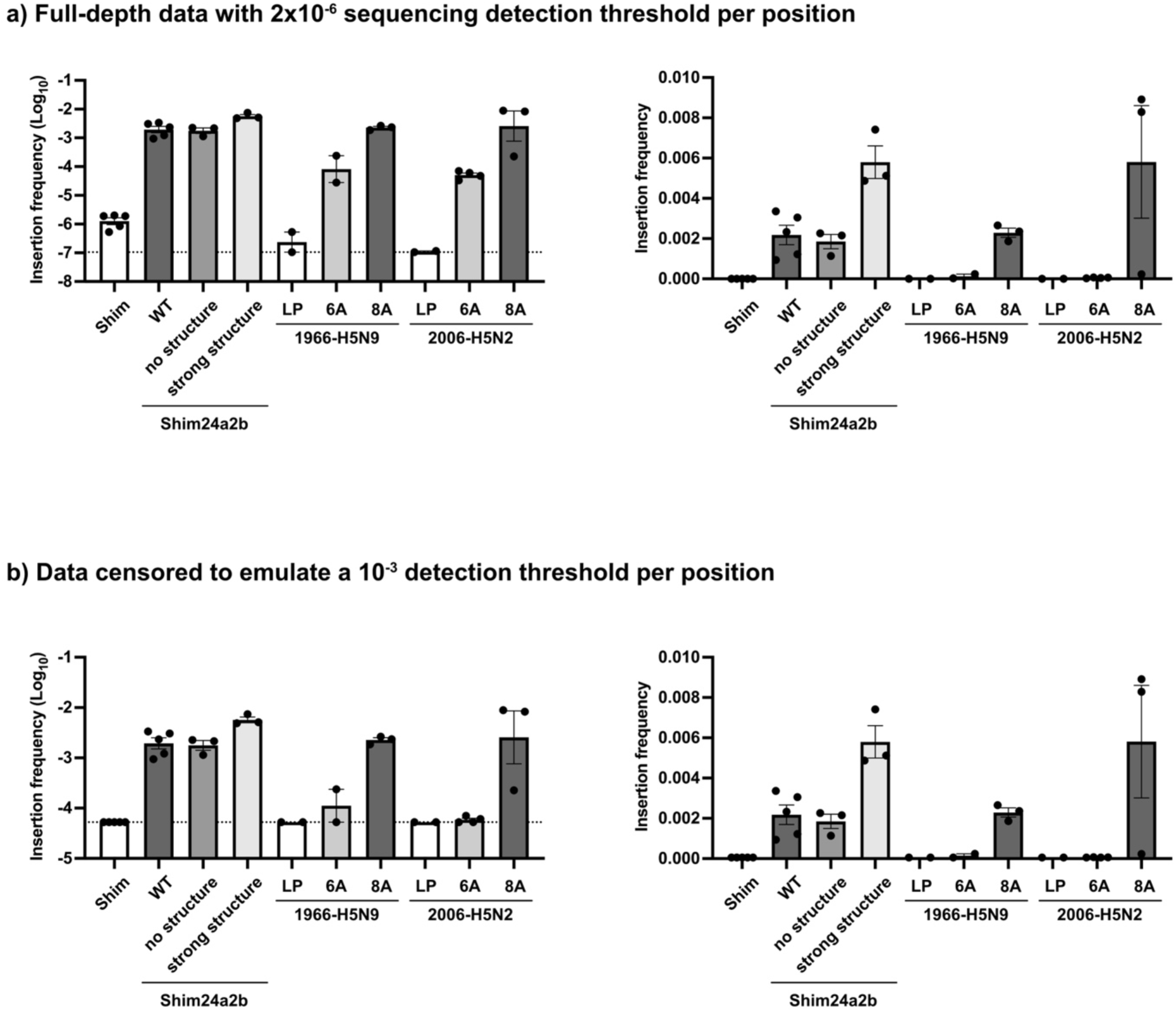
Detection threshold and graphical scaling influence visualization of insertion-frequency distribution. Insertion frequencies were averaged across the 19-nucleotide HA cleavage-site window for PA-H5-Shim, PA-H5-Shim24a2b, its variants presenting different predicted transient structure stabilities and two selected natural H5 HPAIV emergence-event variants. Left panels show insertion frequencies on a Log_10_ scale, whereas right panels show the same data on a linear scale. a) Full-depth dataset, corresponding to a sequencing detection threshold of 2×10-6 per position. Dashed lines indicate the corresponding detection threshold of 1.05×10-7 after averaging across the 19-nucleotide analysed window. b) Data censored to emulate a higher sequencing detection threshold of 10⁻³ per position. Dashed lines indicate the corresponding detection threshold of 5.26×10-5 after averaging across the 19-nucleotide analysed window. Log_10_-scale representation captures insertion frequencies across several orders of magnitude, whereas linear-scale representation emphasizes differences in the higher-frequency range. Censoring to emulate a higher detection threshold further masks lower-frequency insertion events, illustrating how limited dynamic range and linear visualization can accentuate modest differences among high-frequency samples. Bars show mean ± s.e.m.; points represent independent experiments.

**Table S1.**
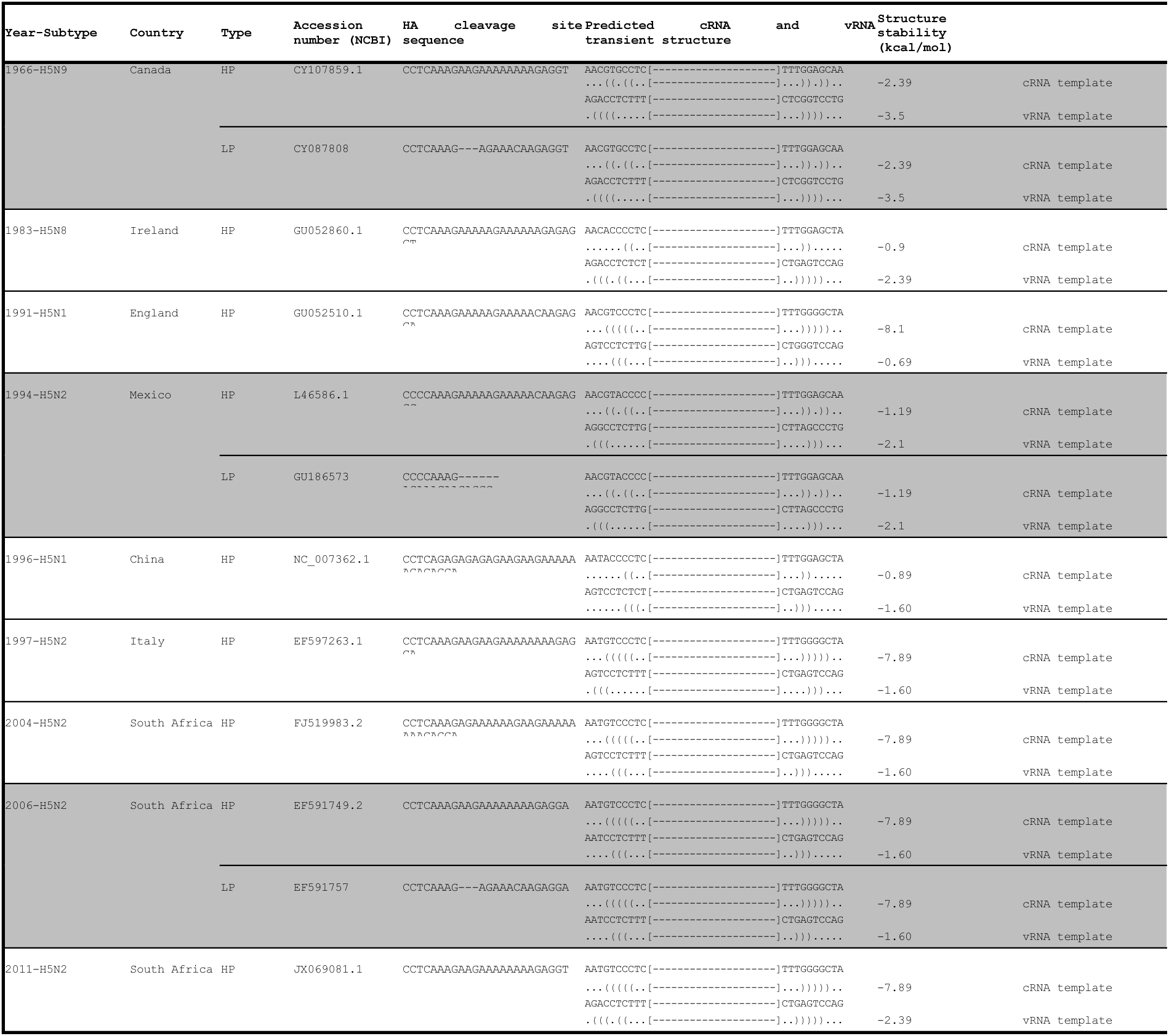
List of insertion-driven H5 HPAIV emergence events. For each HPAIV emergence event, the HPAIV HA cleavage site sequence is provided, along with the predicted cRNA and vRNA transient RNA structure from the inferred precursor for which the MBCS was replaced with a typical H5 LPAIV RETR-encoding sequence. For the well-characterized 1966 H5N9, 1994 H5N2 and 2006 H5N2 emergence events, the LPAIV precursors were identified through co-isolation with HPAIV from the same or neighbouring farms (highlighted in grey). Predicted cRNA and vRNA secondary RNA structures are indicated in dot-bracket format with periods indicating unpaired nucleotides, and brackets indicating G:C, A:U and G:U pairs. Square brackets and hyphens indicate the viral RdRp footprint.

**Table S2.**
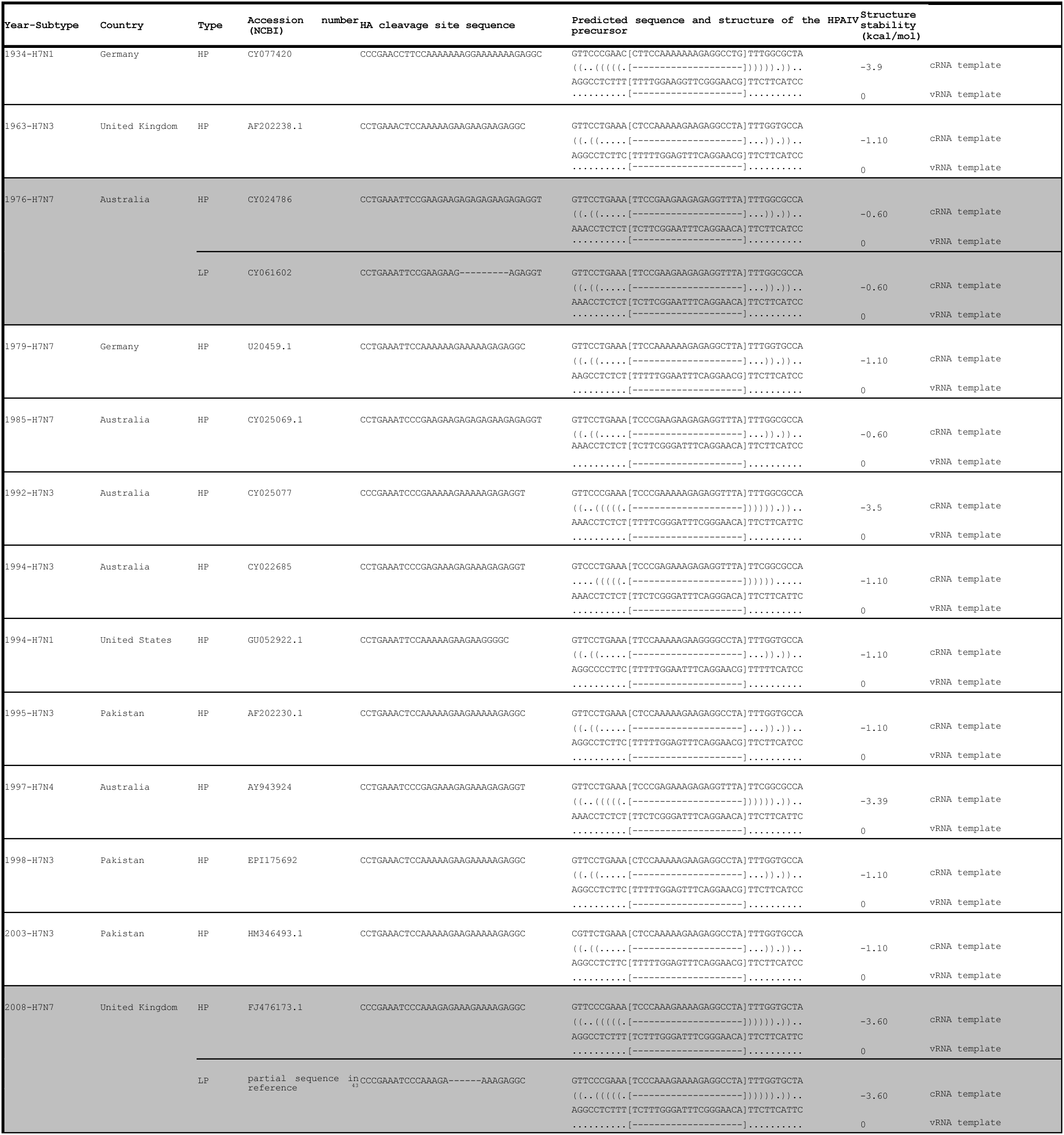
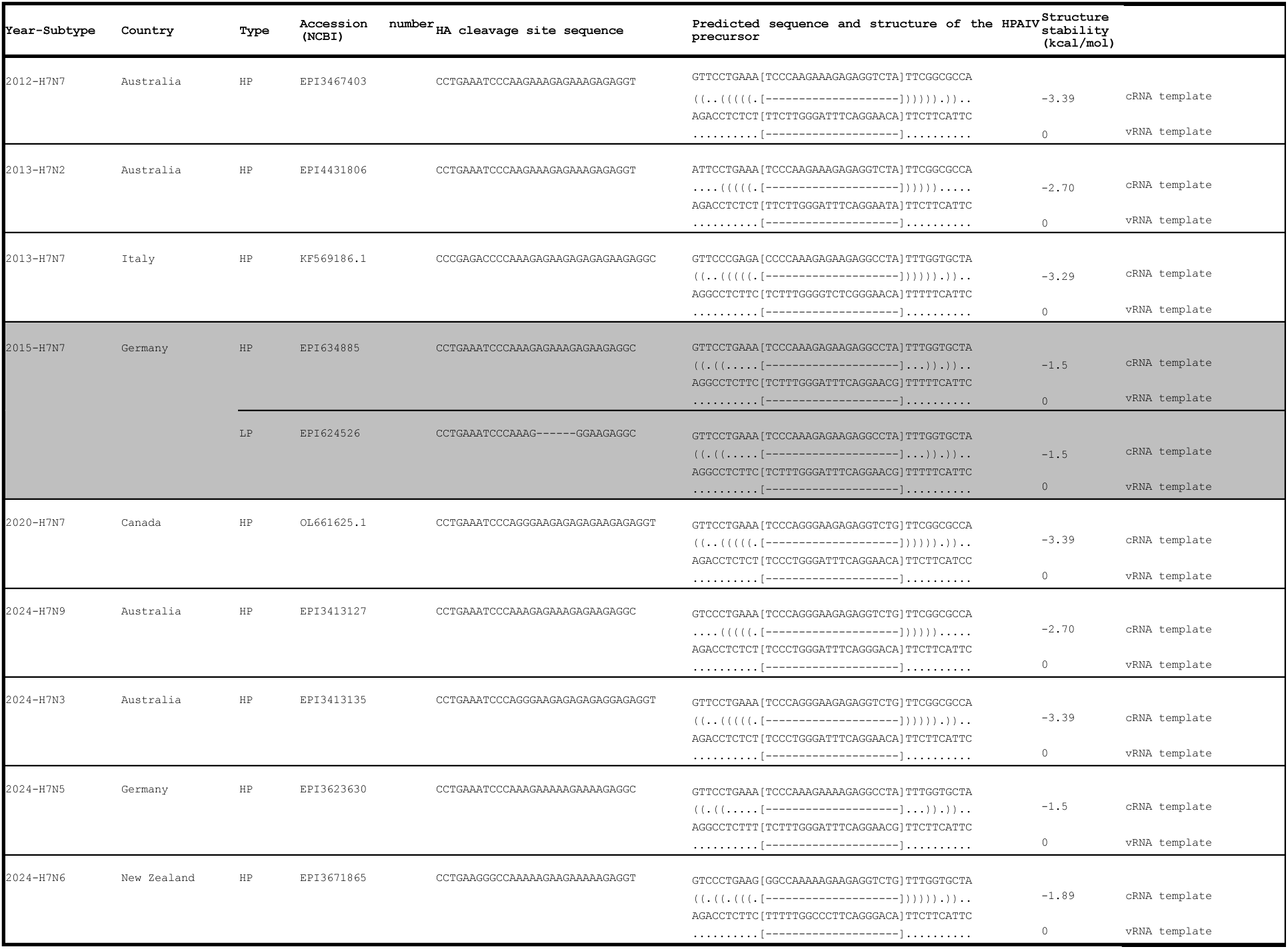
List of insertion-driven H7 HPAIV emergence events. For each HPAIV emergence event, the HPAIV HA cleavage site sequence is provided, along with the predicted cRNA and vRNA transient RNA structure from the inferred precursor for which the MBCS was replaced with a typical H7 LPAIV encoding sequence. For the well-characterized 1976 H7N7, 2008 H7N7 and 2015 H7N7 emergence events, the LPAIV precursors were identified through co-isolation with HPAIV from the same or neighbouring farms (highlighted in grey). Predicted cRNA and vRNA secondary RNA structures are indicated in dot-bracket format with periods indicating unpaired nucleotides, and brackets indicating G:C, A:U and G:U pairs. Square brackets and hyphens indicate the viral RdRp footprint.

**Table S3.** ΔG and ΔΔG thresholds used to classify thermodynamic pathway accessibility.

| Displacement length | Maximum $\Delta G$ threshold values | Maximum $\Delta\Delta G$ threshold values |
| --- | --- | --- |
| Short (<4 nucleotides) | 9 | 15.5 |
| Long ($\geq 4$ nucleotides) | 2.5 | 7.4 |

All values are expressed in kcal mol<sup>-1</sup>. Candidate slippage trajectories were classified as thermodynamically accessible only when all intermediate product–template duplexes along the trajectory satisfied both the $\Delta G$ and $\Delta\Delta G$ thresholds for the corresponding displacement class.

**Data S1. (separate file: Supplementary data 1.xlsx)**

This file contains predicted nucleotide insertion profiles for the H5 dataset, the resulting amino acid sequences, and their predicted furin-cleavability scores.

**Data S2. (separate file: Supplementary data 2.xlsx)**

This file contains predicted nucleotide insertion profiles for the H7 dataset, the resulting amino acid sequences, and their predicted furin-cleavability scores.

**Data S3. (separate file: Supplementary data 3.xlsx)**

This file contains individual SRA run accessions, sample identifiers and associated metadata.

**Data S4. (separate file: Supplementary data 4.xlsx)**

This file contains processed data supporting the figures and analyses.

